# pixOL: pixel-wise dipole-spread function engineering for simultaneously measuring the 3D orientation and 3D localization of dipole-like emitters

**DOI:** 10.1101/2021.12.30.474544

**Authors:** Tingting Wu, Jin Lu, Matthew D. Lew

## Abstract

Interactions between biomolecules are characterized by both where they occur and how they are organized, e.g., the alignment of lipid molecules to form a membrane. However, spatial and angular information are mixed within the image of a fluorescent molecule–the microscope’s dipole-spread function (DSF). We demonstrate the pixOL algorithm for simultaneously optimizing all pixels within a phase mask to produce an engineered Green’s tensor–the dipole extension of point-spread function engineering. The pixOL DSF achieves optimal precision for measuring simultaneously the 3D orientation and 3D location of a single molecule, i.e., 1.14^*°*^ orientation, 0.24 sr wobble angle, 8.17 nm lateral localization, and 12.21 nm axial localization precisions over an 800-nm depth range using 2500 detected photons. The pixOL microscope accurately and precisely resolves the 3D positions and 3D orientations of Nile red within a spherical supported lipid bilayer, resolving both membrane defects and differences in cholesterol concentration, in 6 dimensions.

## INTRODUCTION

The translational and rotational movements of molecules underlie almost all biological and chemical processes. For example, cell membranes are characterized by the organization and alignment of their lipid constituents; the folding conformation or structural disorder of a protein largely determines its interactions with neighbors; DNA must be unwound and accessible by an RNA polymerase for a gene to be expressed. Thus, to study biological function and dysfunction, the positions and conformations of biomolecules are both quantified by molecular dynamics simulations and visualized by experimental imaging techniques. However, imaging these dynamics in native biological environments is difficult. Electron microscopy has exquisite resolution, but cannot be used to image living cells [1, 2]. Interferometric optical scattering can detect, image, and even measure the mass of single molecules (SMs) [3], but it is difficult to distinguish the scattering of one molecular species from another. Superresolution microscopy can now achieve molecular resolution [4–6], but these techniques often intentionally reduce orientation sensitivity (e.g., MINFLUX) so that their position measurements are more robust. Thus, visualizing the position and orientation of SMs simultaneously, precisely, and robustly is difficult. The imaging task is inherently multidimensional, with 6 dimensions of information (3D position and 3D orientation) being conveyed by only hundreds or a few thousand fluorescence photons. Quantifying fundamental limits and engineering optimal methods for maximum measurement precision are topics of active research [7–10].

Many methods have been proposed for measuring either the 3D positions [11] or the 3D orientations [12–14] of SMs. However, there are comparatively few methods experimentally demonstrated for measuring 3D position and 3D orientation simultaneously in single-molecule orientation-localization microscopy (SMOLM). The double-helix PSF [15] and bisected pupil [16] are early examples of PSFs designed for 3D single-molecule localization microscopy (SMLM), but both require relatively bright emitters. More recently, CHIDO uses a stress-engineered (birefringent) optic and polarizing beamsplitter (PBS) in the fluorescence detection path to measure 3D orientations and positions [17]. However, its measurement precision is strongly affected by optical aberrations. In addition, the vortex PSF measures SM 3D position and orientation by modulating fluorescence emission with a vortex phase plate, common in STED nanoscopy, and does not require a PBS [18]. However, its simple implementation requires comparatively bright emitters [13] or a PBS [19] to achieve high orientation measurement precision. Recent reports using estimation theory show that the aforementioned methods do not yet achieve optimal performance [13]. We hypothesize that more powerful optimization tools are required to explore the expansive space of possible Green’s tensors; dipole-spread function (DSF) engineering is distinguished from PSF engineering by the need to optimize for both 3D position and 3D orientation measurement precision–an inherently more complex task. We are inspired by recent methods that leverage neural networks to engineer PSFs with excellent performance for dense 3D SMLM [20].

In this Article, we demonstrate the pixOL microscope, which achieves superior performance over state-of-the-art DSFs for simultaneously measuring the 3D orientations and 3D locations of SMs across an extended depth range (1 µm). We design an algorithm (pixOL) to simultaneously optimize all pixels of a phase mask to shape the dipole response from the microscope, i.e., engineer its Green’s tensor. Unlike optimization using Zernike polynomials [21], pixOL can directly take advantage of supercritical fluorescence arising from imaging SMs near a refractive index interface [22]. The resulting pixOL DSF measures simultaneously the 3D orientations and 3D positions of Nile red (NR) molecules transiently attached to spherical supported lipid bilayers (SLBs). In experiments, SMOLM using the pixOL DSF accurately resolves the 3D spherical shape of the lipid membrane. Further, SMOLM reconstructions show the presence or absence of cholesterol within the membrane through accurate orientation imaging of NR relative to the membrane surface. To our knowledge, these experiments are the first demonstrations of nanoscale super-resolved imaging with accurate molecular 3D position and 3D orientation determination over an entire extended object.

## PIXEL-WISE OPTIMIZATION FOR SMOLM DSF DESIGN

We model a fluorescent molecule as a dipole-like emitter [23–25] with a mean orientation [*µ*_*x*_, *µ*_*y*_, *µ*_*z*_] = [sin *θ* cos *ϕ*, sin *θ* sin *ϕ*, cos *θ*] and a “wobble” solid angle Ω that characterizes its rotational diffusion [26, 27] during a camera frame (Fig. 1(b)). The image produced by the microscope is linearly proportional to a molecule’s orientational second-moment vector 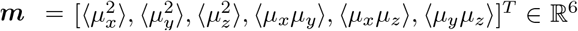, given by (Eqn. (S3))

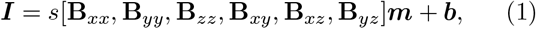

where ***I* ∈** ℝ^*N*^ is the captured intensity on a camera with *N* pixels, *s* is the number of signal photons detected from the emitter, and ***b*** is the background in each pixel. The angle brackets ⟨·⟩ represent a temporal average over one camera frame. The matrices **B**_*il*_ ∈ ℝ^*N*×1^ correspond to the imaging system’s response to each orientational second moment and can be calculated using vectorial diffraction theory [28, 29]. The DSF of any SM is a linear combination of these basis images, which are directly related to the Green’s tensor [27, 30].

**FIG. 1.**
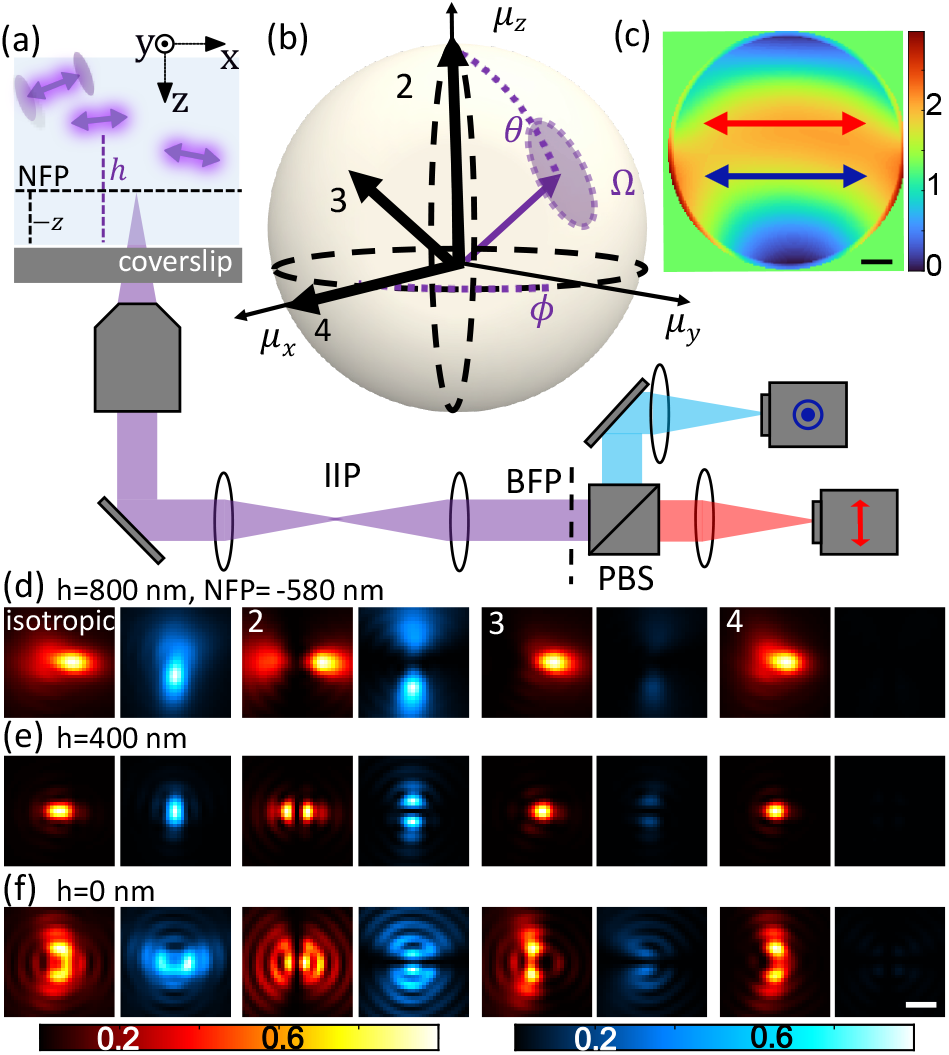
pixOL phase mask and dipole spread function (DSF) for measuring the 3D orientations and 3D positions simultaneously of dipole-like emitters. (a) Imaging system schematic. A microscope objective is focused at a nominal focal plane (NFP, dotted black line) within water at a distance *−z* above (+z below) the coverslip at *z* = 0. The objective collects fluorescence photons from emitters at various locations (*x,y,h*), where *h >* 0 for an emitter above the coverslip. A polarization-sensitive 4f system, comprising 3 lenses and a polarizing beam splitter (PBS), is added after the microscope’s intermediate image plane (IIP) to place the pixOL phase mask at the back focal plane (BFP). Two cameras (or two regions of a single camera) capture x-polarized (red) and y-polarized (cyan) fluorescence. Arrows denote polarization of the light in each channel. (b) Orientation of a dipole-like emitter, parameterized by a polar angle *θ* [0^*°*^, 90^*°*^], azimuthal angle *ϕ* ∈ (*−*180^*°*^, 180^*°*^], and wobble solid angle Ω ∈ [0, 2*π*] sr. (c) Optimized pixOL phase mask. Arrows denote polarization of light in (a) relative to the phase mask. Colormap: phase (rad). Scalebar: 500 µm. (d-f) Simulated images of emitters located at (d) *h* = 800 nm, (e) *h* = 400 nm, and (f) *h* = 0 nm with orientations (*θ, ϕ*, Ω) shown in (b) (emitter 1: Ω = 2*π* sr, emitter 2: (0^*°*^, 0^*°*^, 0), emitter 3: (45^*°*^, 0^*°*^, 0), emitter 4: (90^*°*^, 0^*°*^, 0)), captured in the two polarization channels shown in (a) with the NFP at *z* = *−*580 nm. The intensities of each red-blue image pair are normalized. Scalebar: 500 nm.

A microscope directly encodes a dipole emitter’s lateral position into the location of its shift-invariant DSF, while an SM’s axial location (*h*) and 3D orientation (*θ, ϕ*, Ω) are hidden in the shape of the DSF. To achieve high precision for estimating 3D orientation and 3D location, the shape of the DSF must vary quickly as an emitter’s orientation and axial location changes. This measurement sensitivity can be quantified using the Fisher information matrix [31]. Its matrix inverse, the Cramér-Rao bound (CRB), gives a lower bound on the variance of any unbiased estimator.

We use the CRB matrix **K**, which quantifies the performance of estimating the orientational second moments ***m***, to optimize a phase mask for a polarization-sensitive microscope (Fig. 1(a), SI Section 1). We specifically consider emitters that are located near a water-glass interface, which enables super-critical fluorescence to be captured by the imaging system, thereby boosting measurement sensitivity [22]. To best leverage this information, we simultaneously optimize all pixels of a phase mask ***P*** ∈ ℝ^*n×n*^ by minimizing the loss function

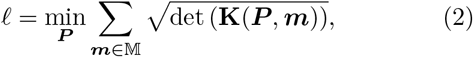

where 𝕄 denotes orientation space (SI Section 1) and det(·) represents the determinant of a matrix. To avoid poor lateral estimation precision, we force the algorithm to create DSFs smaller than 1.8 *µ*m × 1.8 *µ*m by ignoring photons diffracted outside of this region on the camera.

The optimized pixOL phase mask (Fig. 1(c)) breaks the symmetries within the images produced by the six orientational moments ***m*** at the back focal plane (Fig. S1). Notably, it modulates super-critical fluorescence differently from the rest of the BFP, indicating that the algorithm intelligently optimizes the response of individual pixels to maximize orientation measurement sensitivity (Fig. S1). The resulting DSFs of SMs of various orientations exhibit easily discernible shapes and intensities across the x- and y-polarized imaging channels (Fig. 1(d-f)), owing to their 6 distinct basis images (Fig. S3). Simultaneously, pixOL also breaks the symmetry of defocus, rotating the DSF by 90^*°*^ when an SM is above vs. below the focal plane (Fig. 1(d-f)).

Using CRB as a performance metric, we compare our pixOL DSF to other engineered DSFs designed for 3D orientation and 3D position measurements, namely the double helix [15], CHIDO [17], and unpolarized vortex DSFs [18] (SI Section 4, see Fig. S5for additional comparisons to DSFs only designed for orientation). We calculate the mean angular standard deviation *σ*_*δ*_ (MASD, Eqn. S23) as a combined precision of measuring 3D orientation (*θ, ϕ*) and the standard deviation *σ*_Ω_ of measuring wobble. For in-focus (Fig. 2(a,b)) emitters, pixOL shows the best precision for measuring 3D orientation (mean *σ*_*δ*_ = 0.80^*°*^, 10% better MASD than the next-best DSF, CHIDO, and mean *σ*_Ω_ = 0.16 sr, 11% better wobble precision than CHIDO). Over an 800-nm depth range, pixOL’s orientation precision degrades slightly (mean *σ*_*δ*_ = 1.14^*°*^ and mean *σ*_Ω_ = 0.24 sr) and is comparable to CHIDO (Fig. S6). We also quantified the lateral localization precision *σ*_*L*_ and the axial localization precision *σ*_*h*_ for isotropic emitters across an axial range of 800 nm (Fig. 2(c,d)). The pixOL DSF has superior lateral precision compared to all other DSFs (mean *σ*_*L*_ = 8.17 nm over an 800-nm depth range, 26% better than CHIDO). It also has excellent axial localization precision (mean *σ*_*h*_ = 12.21 nm over an 800-nm depth range), outperformed only by the double-helix.

**FIG. 2.**
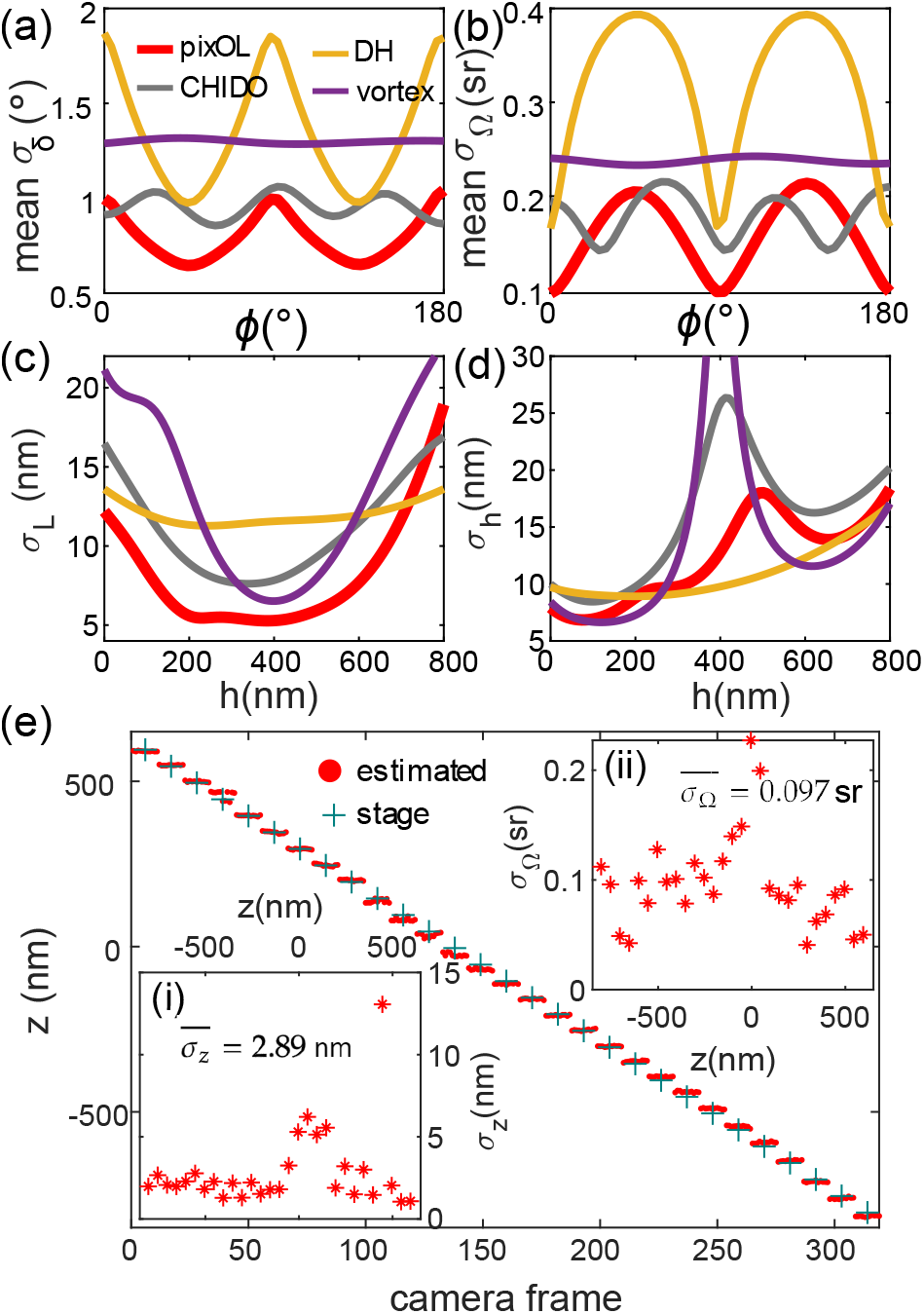
Measurement precision of the pixOL DSF. (a-d) Best-possible measurement precision of various engineered DSFs calculated using the Cramér-Rao bound (CRB) for emitters within water (1.33 refractive index) with 2500 signal photons and 3 background photons per pixel detected. Red: pixOL, yellow: double helix (DH) [15], grey: CHIDO [17], purple: unpolarized vortex [18]. (a) Mean angular standard deviation *σ*_*δ*_ (MASD) averaged uniformly over all *θ*. MASD quantifies the combined precision of measuring *θ* and *ϕ* as the half-angle of a cone representing orientation uncertainty (Eqn. S23) [32]. (b) Mean wobble angle precision *σ*_Ω_ averaged uniformly over all *θ*. (c,d) Localization precisions *σ*_*L*_ and *σ*_*h*_ for measuring (c) lateral position *l* and (d) axial location *h* above the interface, respectively. MASD and *σ*_Ω_ are calculated for in-focus SMs with fixed orientation (Ω = 0 sr); localization precisions are for isotropic emitters (Ω = 2*π* sr). (e) Position and emission anisotropy measurements of a fluorescent bead, scanned axially from *z* = −790 nm to *z* = 610 nm with a step size of 50 nm (11 camera frames per step). Red dot: estimated axial distance *z* between the bead and focal plane in each frame; green cross: expected stage position. Inset (i): Experimental axial precision *σ*_*z*_ at each scanning plane (mean precision 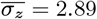 nm). Inset (ii): Experimental emission anisotropy precision *σ*_Ω_ at each scanning plane (average precision 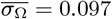 sr).

In any experiment, optical aberrations will perturb the designed DSF and decrease estimation performance. Using a liquid-crystal spatial light modulator placed in the microscope back focal plane (Fig. 1), we compare the DSF produced by the pixOL phase mask (Fig. S7) to that of its conjugate (pixOL*, Fig. S8). Interestingly, the pixOL* DSF better matches the depth-dependent features of the ideal pixOL DSF. Thus, we use the pixOL* mask, an experimentally calibrated DSF model (SI Section 3), and a bespoke regularized maximum-likelihood estimator (SI Section 2) to jointly estimate the 3D orientations and 3D positions of all SMs within each FOV. Monte Carlo simulations of our estimation algorithm show that the pixOL microscope achieves 23.2 nm lateral and 19.5 nm axial localization precisions on average throughout a 700-nm depth range with 2500 detected photons (Figs. S9, S10, and S11). Simultaneously, pixOL estimates the orientation and wobble of each SM with 4.1^*°*^ and 0.44 sr precisions, respectively.

Scanning fluorescent beads (100-nm diameter) across an axial range of 1400 nm enables us to verify localization and orientation measurement precisions experimentally. The trajectory of defocus estimates *z* resolves the 50-nm stage movements very well (Fig. 2(e), average axial precision 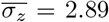 nm in Fig. 2(e)(i)). Since the bead contains many fluorophores, we may quantify the bead’s emission pattern by measuring its effective “wobble” angle Ω. We find that its emission is largely isotropic (wobble angle 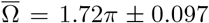 sr averaged over all steps, mean±std, Figs. 2(e)(ii) and S14(e)). In our implementation, experimental precisions for measuring axial location (Fig. 2(e)(i)) and orientation (Fig. 2(e)(ii)) can degrade slightly for certain emitter orientations when they are in focus (Fig. S10); this degradation can be corrected with additional aberration corrections. Data from other beads also show precise localization and orientation estimates (Fig. S14).

## 6D SMOLM REVEALS MEMBRANE MORPHOLOGY AND COMPOSITION

To demonstrate accurate 6D imaging of molecular orientations (*θ, ϕ*, Ω) and positions (*x, y, h*), we adhere supported lipid bilayers (SLBs) to silica beads (2 *µ*m diameter, Fig. 3(a), SI Section 6) [33, 34] using two lipid compositions: DPPC (di(16:0) phosphatidylcholine) only vs. a mixture of DPPC and cholesterol (chol). Previous studies [19, 35] have shown that NR orients itself perpendicular to the membrane when a high concentration (40% used here) of chol is present. Thus, a spherical SLB enables us to validate simultaneously orientation and position imaging performance, quantifying both precision and accuracy. We use the transient binding and blinking of Nile red (NR) molecules to the SLBs [35–37] to facilitate SM detection and orientation-position measurements (Fig. 3(a), Movie S1). The beads are illuminated by a tilted, circularly polarized laser beam so that emitters can be excited efficiently regardless of their 3D orientation.

**FIG. 3.**
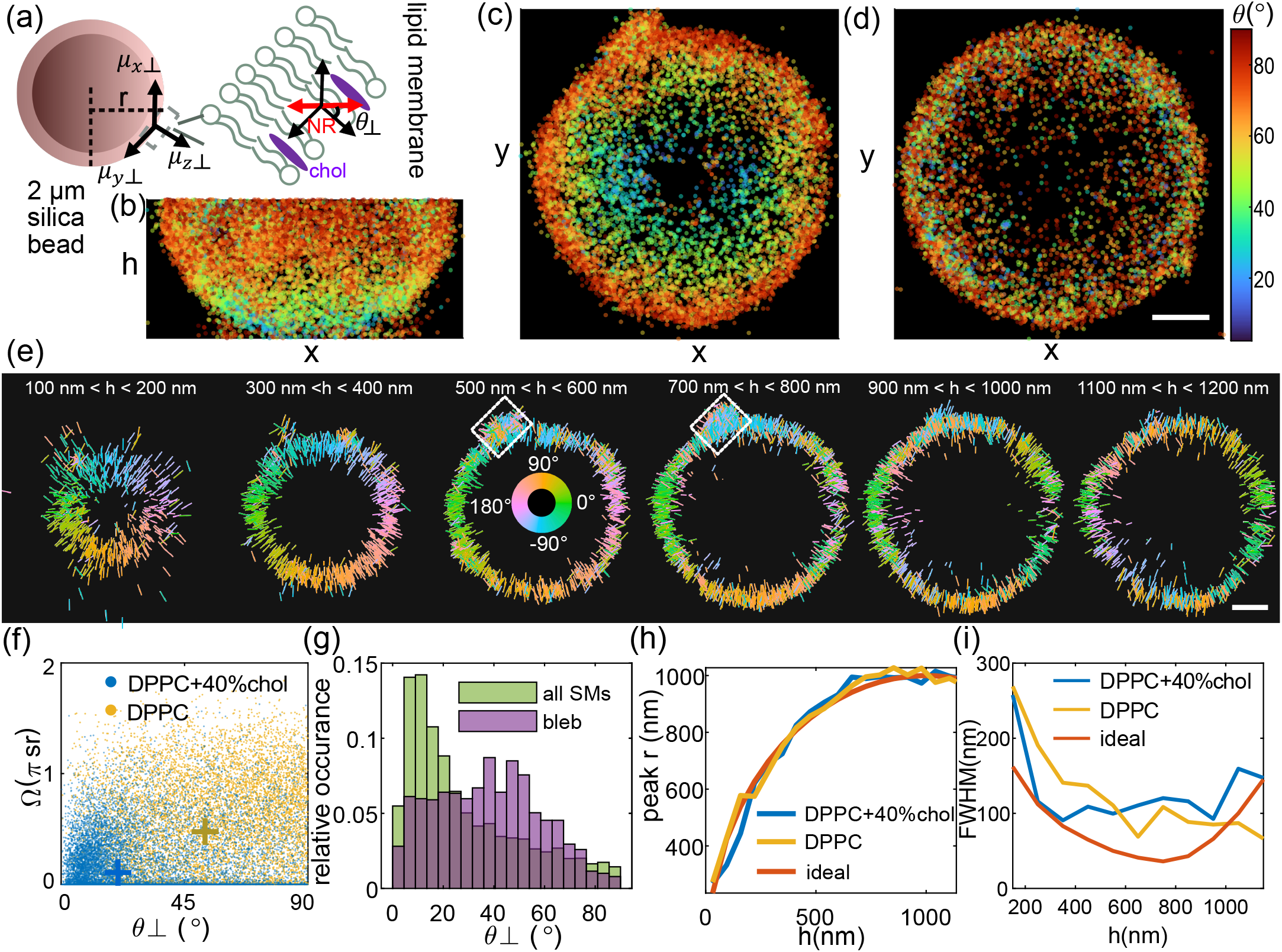
SMOLM images of the 3D orientations and 3D locations of Nile red (NR) within spherical supported lipid bilayers (SLBs) collected by the pixOL DSF. (a) SLBs are adhered to 2 µm-diameter silica spheres, where *θ*_⊥_ represents the relative angle between the orientation of each NR molecule and the SLB’s surface normal and *r* is the distance between the 2D location (x,y) of a NR within a certain *h* slice and the center of the sphere. (b-d) 2D maps of polar orientation *θ* for each NR molecule, shown for the bottom half of each bead. (b) x-h and (c) x-y views for an SLB consisting of DPPC and 40% cholesterol (chol). (d) x-y view of a DPPC-only SLB. Colorbar: *θ* (deg). (e) x-y cross-sections of the bead in (b,c) depicting the 3D orientation (*θ,ϕ*) of each NR as a line segment. The length and direction of each line indicate the magnitude 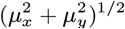 and direction *ϕ* of azimuthal orientation. Colors represent azimuthal orientation *ϕ*. White box: a membrane bleb. (f) NR orientation *θ*_⊥_ vs. wobble Ω for the (blue) DPPC+chol SLB and (yellow) DPPC-only SLB. Crosses indicate measurement medians. (g) NR orientations *θ*_⊥_ (green) across the entire DPPC+chol SLB and (purple) within the membrane bleb in (e). (h,i) Measured (h) cross-sectional radius *r* and (i) apparent thickness of the spherical SLB, calculated as the peak and full-width at half-maximum (FWHM), respectively, of the distribution of NR lateral positions *r*, in each *h* slice in (e) and Figs. S16(d), S17(e). (blue) DPPC + chol bead, (yellow) DPPC-only bead, and (red) ideal sphere. The theoretical FWHM in (i) accounts for the projection of the SLB into the xy plane and pixOL’s localization precision (Eqn. S27). Scale bars: 400 nm.

For the SLB containing chol, the 3D locations of NR form a sphere as expected (Fig. S16(a)). The orientations of NR change smoothly from being mostly parallel to the optical axis (small *θ*) at the bottom of the sphere to being within the xy-plane (*θ* approaching 90^*°*^) at the sphere’s waist (Fig. 3(b,c), Movie S2). Likewise, the azimuthal orientations *ϕ* of NR show that each molecule is perpendicular to the sphere’s surface within each *h* slice (Fig. 3(e), Fig. S16(a,b), Movie S3). Note that the sphere’s shape and the symmetry of SM emission guarantee exactly two locations, on opposite sides of the sphere, where NR orientations are identical to one another. For example, the measured *ϕ* below the sphere’s equator (*h <* 900 nm) match the *ϕ* measurements on the opposite surface above the equator (*h >* 1100 nm, Fig. 3(e)). Calculating the relative orientation *θ*_⊥_ between each NR and the surface normal of the sphere shows that the molecules lie mostly parallel (small *θ*_⊥_) to the lipid tails within the SLB (Fig. 3(f)), regardless of their location on the sphere (Fig. S16(c)). Moreover, each NR exhibits relatively little rotational diffusion, i.e., small wobble angle Ω, (Fig. 3(f)), which is consistent with previous characterizations of NR within planar SLBs [19, 35].

Interestingly, we detect a bleb in the membrane (white boxes in Fig. 3(e)) where NR orientations *θ*_⊥_ are more varied (Fig. 3(g)), showing the bleb’s local disorganization. However, NR wobbles Ω within the bleb are similar to other regions of the sphere (Fig. S16(f)), implying that chol is distributed uniformly throughout the membrane. Without cholesterol, DPPC molecules within the SLB exhibit greater intermolecular spacing. NR in contact with the DPPC-coated bead reveals the absence of cholesterol via more random orientations *θ, ϕ*, and *θ*_⊥_ (Fig. 3(d,f), Fig. S17(a-d), Movies S2, S3) than those of the SLB containing cholesterol (median 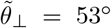 for DPPC only vs. 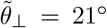 for DPPC+chol). NR also shows larger wobble angles (median 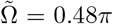 sr for DPPC only vs. 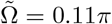 sr for DPPC+chol, Fig. 3(f))–another indication of less crowding within the pure DPPC SLB.

We quantify the shape and apparent thickness of the SLB by calculating the (2D) radial distance *r* between each NR location and the sphere’s center within each *h* slice (Figs. S16(d), S17(e)). The estimated shapes of two beads match the expected cross-sectional radius of an ideal sphere accurately (Fig. 3(h)). We also compute the best-possible full-width at half-maximum (FWHM) that the pixOL* microscope can achieve, accounting for the curvature of the spherical surface and assuming that pixOL* achieves CRB-limited localization precision (SI Section 7). On average, the apparent SLB thickness measured by pixOL* is 55% larger than the best-possible precision of pixOL* (Eqn. S27) across a 1200 nm axial range (average FWHM is 129 nm for DPPC with cholesterol, 123 nm for DPPC only, and 82 nm for the theoretical distribution, Fig. 3(i)). This distribution is significantly broader than the CRB and likely stems from optical aberrations (Fig. S8) and precision and bias from our estimation algorithm (Fig. S9-S11). Estimation performance can be improved via more detailed aberration calibrations and corrections [18], as well as more powerful estimation algorithms that robustly explore 6-dimensional position-orientation space with greater accuracy and computational efficiency (SI Section 5).

## CONCLUSION

Here, we propose an algorithm (pixOL) for dipole-spread function (DSF) engineering, i.e., using vectorial diffraction theory to simultaneously optimize all pixels of a phase mask for measuring the 3D orientation of dipole-like emitters. The resulting pixOL DSF achieves superior orientation and localization precision across a larger axial range than other techniques (Fig. 2). Pixel-wise engineering of its phase mask enables the pixOL DSF to leverage super-critical fluorescence (Fig. 1(c)) to improve orientation-measurement sensitivity [22]. Super-critical fluorescence is also beneficial for improving the pixOL DSF’s axial localization precision, as utilized by DONALD [38] and DAISY [39], since the amount of supercritical light can be used to measure *h*, the height of an emitter above an refractive index interface. In addition, one can easily modify pixOL’s imaging model to generate phase masks that are optimal for other imaging geometries.

Notably, the use of DNA PAINT and DNA origami as molecular “rulers” is the gold standard for validating the accuracy of optical nanoscopic tools [40, 41]. However, due to practical issues with the robustness and precision of controlling both the 3D positions and 3D orientations of the labels in these samples, we adapted 3D spherical SLBs [33, 34] to demonstrate experimentally the accuracy and precision of the pixOL microscope for 6D SMOLM imaging. Visualizing SMOLM data of NR binding to these SLBs shows highly spherical membrane morphologies and dye orientations that are perpendicular to the spherical surface (Fig. 3). Thus, despite the presence of aberrations typical in optical microscopes, the diverse features of the pixOL DSF are detectable in the images of flashing NR (Fig. S16(e)) and convey the 3D orientations and 3D positions of fluorescent molecules accurately and precisely. Moreover, our imaging of the spherical SLBs shows that the pixOL DSF sensitively discerns membrane morphology and the composition of its lipid components through detailed measurements of NR positions, orientations, and rotational diffusion.

Since the pixOL microscope measures the 3D orientations and 3D locations simultaneously of SMs, we anticipate the technology enabling fascinating studies of biomolecular interactions away from the coverslip, e.g., the 3D growth of amyloid aggregates and their interactions with cellular membranes. However, additional developments can further improve SMOLM’s versatility for biological studies. In addition to improving position and orientation measurement precision, DSFs capable of coping with high background autofluorescence, as is typical in cellular imaging [14], as well as advanced machine learning algorithms [20] able to distinguish and localize molecules whose images overlap on the camera, are needed. These topics, as well as the development of DSFs whose performance is closer to fundamental limits [8–10], remain exciting directions for future research.

## Acknowledgments

The authors thank Oumeng Zhang and Tianben Ding for helpful suggestions and comments.

## Disclosures

The authors declare no conflicts of interest.

See Supplement 1 for supporting content. The pixOL algorithm, estimation algorithm, and data of spherical supported lipid bilayers are available via OSF [42] and by request.

