## Supplemental material 1 for "pixOL: pixel-wise dipole-spread function engineering for simultaneously measuring the 3D orientation and 3D localization of dipole-like emitters"

This document provides supplementary information to "pixOL: pixel-wise dipole-spread function engineering for simultaneously measuring the 3D orientation and 3D localization of dipole-like emitters," offering details on the optimization algorithm, estimation algorithm, experimental calibration, and sample preparation.

**CONTENTS**

**List of Figures** 1

1

**I Imaging model and pixOL algorithm** 2

**II Single-molecule detection and 3D orientation and 3D position estimation** 2

**III Imaging system calibration** 5

**IV Quantifying the precision of measuring 3D orientation and 3D position** 5

**V Quantifying estimation bias when measuring 3D orientation and 3D position** 6

**VI Preparing 3D spherical supported lipid bilayers** 6

**VII Imaging spherical lipid bilayers using Nile red** 6

**References** 19

*

### I. IMAGING MODEL AND PIXOL ALGORITHM

The goal of the pixOL algorithm is to simultaneously optimize all pixels of a phase mask $\boldsymbol{P} \in \mathbb{R}^{n\times n}$ so that the resulting dipole-spread function (DSF) has optimal precision for measuring the 3D orientation $[\mu_x, \mu_y, \mu_z, \Omega]$ of dipole-like emitters. The dipole's average orientation $(\theta, \phi)$ in spherical coordinates during a camera frame is related to its unit vector parameterization $\boldsymbol{\mu} = [\mu_x, \mu_y, \mu_z]^T$ in Cartesian space by

$$[\mu_x, \mu_y, \mu_z] = [\sin\theta\cos\phi, \sin\theta\sin\phi, \cos\theta]. \tag{S1}$$

A wobbling dipole with the orientation parameters $[\mu_x, \mu_y, \mu_z, \Omega]$, where the solid angle $\Omega$ characterizes rotational diffusion within a hard-edged cone, may be characterized using the six orientational second moments $\boldsymbol{m} \in \mathbb{R}^6$ as [1, 2]

$$\boldsymbol{m} = \left[\langle\mu_x^2\rangle, \langle\mu_y^2\rangle, \langle\mu_z^2\rangle, \langle\mu_x\mu_y\rangle, \langle\mu_x\mu_z\rangle, \langle\mu_y\mu_z\rangle\right]^T \tag{S2}$$

where

$$\langle\mu_x^2\rangle = \gamma\mu_x^2 + (1-\gamma)/3, \tag{S3a}$$
$$\langle\mu_y^2\rangle = \gamma\mu_y^2 + (1-\gamma)/3, \tag{S3b}$$
$$\langle\mu_z^2\rangle = \gamma\mu_z^2 + (1-\gamma)/3, \tag{S3c}$$
$$\langle\mu_x\mu_y\rangle = \gamma\mu_x\mu_y, \tag{S3d}$$
$$\langle\mu_x\mu_z\rangle = \gamma\mu_x\mu_z, \tag{S3e}$$
$$\langle\mu_y\mu_z\rangle = \gamma\mu_y\mu_z, \tag{S3f}$$
$$\gamma = 1 - \frac{3\Omega}{4\pi} + \frac{\Omega^2}{8\pi^2}. \tag{S3g}$$

We optimize the phase mask for a microscope that splits the collected fluorescence into x- and y-polarized channels (Fig. 1(a)). To model such a system, we begin by writing the x- and y-polarized electric field $\boldsymbol{E}_{\text{BFP}}$ at the back focal plane (BFP) as [1, 3–5]

$$\boldsymbol{E}_{\text{BFP}}(u,v) = \exp\left(jkz_h\sqrt{1-u^2-v^2}\right) \begin{bmatrix} g^{(x)}_{x,\text{BFP}}(u,v) & g^{(x)}_{y,\text{BFP}}(u,v) & g^{(x)}_{z,\text{BFP}}(u,v) \\ g^{(y)}_{x,\text{BFP}}(u,v) & g^{(y)}_{y,\text{BFP}}(u,v) & g^{(y)}_{z,\text{BFP}}(u,v) \end{bmatrix} \begin{bmatrix} \mu_x \\ \mu_y \\ \mu_z \end{bmatrix}, \tag{S4}$$

where $g^{(l)}_{i,\text{BFP}}$ are the basis fields observed in an $l$-polarized BFP from an in-focus ($z_h = 0$) dipole with orientation $\mu_i$ [1, 3–5] and $i, l \in \{x, y, z\}$. The wavenumber $k = 2\pi/\lambda$ represents the wave propagation constant in water, and the BFP coordinates are normalized such that $u^2 + v^2 < 1$.

For computational convenience, we move to a discrete model $\boldsymbol{g}_i$ of the basis fields, sampling the BFP with sufficient pixels and zero-padding to match the spatial light modulator (SLM) of our imaging system. We concatenate the polarized basis fields $[\boldsymbol{g}_i^{(x)}, \boldsymbol{g}_i^{(y)}]^T$ at the image plane of the microscope and express them jointly as

$$\boldsymbol{g}_i = \begin{bmatrix} \boldsymbol{g}_i^{(x)} \\ \boldsymbol{g}_i^{(y)} \end{bmatrix} = \mathcal{F}\left\{ \begin{bmatrix} \exp(j\boldsymbol{P}) \\ \exp(j\boldsymbol{P}) \end{bmatrix} \odot \begin{bmatrix} \boldsymbol{g}^{(x)}_{i,\text{BFP}} \\ \boldsymbol{g}^{(y)}_{i,\text{BFP}} \end{bmatrix} \right\}, \tag{S5}$$

where $\mathcal{F}\{\cdot\}$ denotes a discrete 2D Fourier transform, $\odot$ represents element-wise multiplication of two vectors or matrices. We may now calculate the basis images $\mathbf{B} = [\mathbf{B}_{xx}, \mathbf{B}_{yy}, \mathbf{B}_{zz}, \mathbf{B}_{xy}, \mathbf{B}_{xz}, \mathbf{B}_{yz}] \in \mathbb{R}^{N\times 6}$ in the image plane as

$$\mathbf{B}_{xx} = \boldsymbol{g}_x \odot \boldsymbol{g}_x^* \tag{S6a}$$
$$\mathbf{B}_{yy} = \boldsymbol{g}_y \odot \boldsymbol{g}_y^* \tag{S6b}$$
$$\mathbf{B}_{zz} = \boldsymbol{g}_z \odot \boldsymbol{g}_z^* \tag{S6c}$$
$$\mathbf{B}_{xy} = \boldsymbol{g}_x \odot \boldsymbol{g}_y^* + \boldsymbol{g}_x^* \odot \boldsymbol{g}_y \tag{S6d}$$
$$\mathbf{B}_{xz} = \boldsymbol{g}_x \odot \boldsymbol{g}_z^* + \boldsymbol{g}_x^* \odot \boldsymbol{g}_z \tag{S6e}$$
$$\mathbf{B}_{yz} = \boldsymbol{g}_y \odot \boldsymbol{g}_z^* + \boldsymbol{g}_y^* \odot \boldsymbol{g}_z. \tag{S6f}$$

These images $\mathbf{B}_{il}$ correspond to the DSF (Eqn. 1) produced by a dipole with an orientational second moment $m_{il}$, where $i, l \in \{x, y, z\}$. A similar expression can be derived for the basis images $\mathbf{B}_{\text{BFP}}$ in the back focal plane (Fig. S1).

To calculate the best possible precision for estimating the orientational moments $\boldsymbol{m}$, we calculate the Cramér-Rao bound (CRB) matrix $\mathbf{K}$ using

$$\mathbf{K} = \sum_{j=1}^{N} \frac{s^2}{I_j} \mathbf{B}_j^T \mathbf{B}_j, \tag{S7}$$

where $\mathbf{B}_j \in \mathbb{R}^{1\times 6}$ is the $j^{\text{th}}$ raw of $\mathbf{B}$, the superscript $T$ denotes a matrix transpose. We wish to find an optimal DSF that minimizes $\mathbf{K}$ for any possible orientation $[\mu_x, \mu_y, \mu_z, \Omega]$, so we build our loss function $\ell$ (Eqn. 2) to represent the sum of the precision of second moments over a uniformly sampled orientation space $\mathbb{M}$. The mean orientation $\boldsymbol{\mu} = [\mu_x, \mu_y, \mu_z]^T$ is sampled using [6]

$$\mu_x = 2x_1\sqrt{1-x_1^2-x_2^2} \tag{S8a}$$
$$\mu_y = 2x_2\sqrt{1-x_1^2-x_2^2} \tag{S8b}$$
$$\mu_z = 1 - 2(x_1^2 + x_2^2), \tag{S8c}$$

where $x_1$, $x_2$ are uniformly distributed within $(-1, 1)$ and points for which $x_1^2 + x_2^2 \geq 1$ are rejected. The wobbling angle $\Omega$ is uniformly sampled within $[0, 2\pi]$.

Using GradientTape for automatic differentiation in Tensorflow, we can easily calculate the gradient $\boldsymbol{D}$ of the loss function $\ell$ with respect to the current phase mask $\boldsymbol{P}$. We optimize the mask for dipole emitters located at the glass-water interface with 380 total signal photons detected and 2 background photons in each pixel. To begin, we randomly initialize the phase of each pixel. Updating the current phase mask using the Adam algorithm with a learning rate of 0.05 and a total of 300 iterations, the pixOL algorithm produces a phase mask $\exp(j\boldsymbol{P})$ shown in Fig. 1(c).

### II. SINGLE-MOLECULE DETECTION AND 3D ORIENTATION AND 3D POSITION ESTIMATION

Detecting single molecules (SMs) and estimating their 3D positions and orientations (6 second moments $\boldsymbol{m}$) can be computationally expensive and non-convex. We overcome these challenges by 1) applying a linear approximation to the forward imaging model and 2) separating the detection and estimation process into sequential steps.

To begin, we extend the forward model (Eqn. 1) to accommodate images containing $Q \geq 1$ SMs such that

$$\boldsymbol{I} = \sum_q s_q \mathbf{B}(x_q, y_q, h_q)\boldsymbol{m}_q + \boldsymbol{b}, \tag{S9}$$

where $s_q$ is the brightness in photons, $[x_q, y_q, h_q]$ is the 3D location, and $\boldsymbol{m}_q$ is the orientational second moment vector of the $q^{\text{th}}$ emitter. The basis images $\mathbf{B}$ change shape for SMs at different axial locations, but only shifts linearly for SMs at different lateral positions. To reduce the computational burden of 3D localization, we discretize the continuous 3D location space using a first-order polynomial approximation. Our imaging model becomes

$$\begin{aligned}\boldsymbol{I} = \sum_q \Bigg[ & s_q \mathbf{B}(x_{q0}, y_{q0}, h_{q0})\boldsymbol{m}_q \\ & + s_q \frac{\partial \mathbf{B}(x_{q0}, y_{q0}, h_{q0})}{\partial x}\boldsymbol{m}_q dx_q + s_q \frac{\partial \mathbf{B}(x_{q0}, y_{q0}, h_{q0})}{\partial y}\boldsymbol{m}_q dy_q \\ & + s_q \frac{\partial \mathbf{B}(x_{q0}, y_{q0}, h_{q0})}{\partial h}\boldsymbol{m}_q dz_q \Bigg] + \boldsymbol{b},\end{aligned} \tag{S10}$$

where $[x_{q0}, y_{q0}, h_{q0}]$ is the closest discrete grid point to the continuous location $[x_q, y_q, h_q]$. The off-grid distances $[dx_q, dy_q, dh_q] = [x_q, y_q, h_q] - [x_{q0}, y_{q0}, h_{q0}]$ characterize the difference between the true position and the closest grid point. As the last three columns (i.e., images) of the basis matrix $\mathbf{B}$ have a total energy of zero (Fig. S3), we can further simplify the forward model by excluding these last three images in the first-order approximation. The forward model becomes

$$\boldsymbol{I} = \sum_q \mathbf{A}_q \boldsymbol{\zeta}_q + \boldsymbol{b}, \tag{S11}$$

where

$$\begin{aligned}\mathbf{A}_q = \Bigg[ & \mathbf{B}(x_{q0}, y_{q0}, h_{q0}), \frac{\partial \mathbf{B}^o(x_{q0}, y_{q0}, h_{q0})}{\partial x}, \\ & \frac{\partial \mathbf{B}^o(x_{q0}, y_{q0}, h_{q0})}{\partial y}, \frac{\partial \mathbf{B}^o(x_{k0}, y_{q0}, h_{q0})}{\partial h} \Bigg],\end{aligned} \tag{S12a}$$

$$\boldsymbol{\zeta}_q = \left[ s_q \boldsymbol{m}_q^T, s_q (\boldsymbol{m}_q^o)^T dx_q, s_q (\boldsymbol{m}_q^o)^T dy_q, s_q (\boldsymbol{m}_q^o)^T dh_q \right]^T, \tag{S12b}$$

the basis matrix $\mathbf{B}^o$ excludes the last three basis images, and the second moment vector $\boldsymbol{m}_q^o$ excludes the last three elements. Importantly, the matrix $\mathbf{A}_q$ may be precomputed for a specific imaging system and choice of location grid, while $\boldsymbol{\zeta}_q$ contains the molecular brightness, orientation, and position information we wish to estimate. Thus, we have simplified our 3D orientation and 3D position estimation problem to use a linear forward model (Eqn. S11) involving only 15 elements in $\boldsymbol{\zeta}_q$ for each emitter, i.e., 6 brightness-weighted second moments at each spatial grid point plus 3 brightness- and moment-weighted first-order distances for each off-grid direction. For comparison, the original first-order approximation model in Eqn. S10 involves 24 parameters for each SM.

We perform detection and estimation as separate tasks involving three main parts. We start by determining the number of emitters and their 2D locations within an image. In the second step, we obtain an initial estimate of an emitter's 3D orientation and 3D position based on cropped images centered at each detected SM. Finally, the algorithm updates the estimates of all emitters simultaneously using the entire image.

To simplify the detection process, we ignore the axial dimension and begin with a 2D forward model

$$\boldsymbol{I}^{\text{2D}} = \sum_q \boldsymbol{\zeta}_q^{\text{2D}} \mathbf{A}_q^{\text{2D}} + \boldsymbol{b}, \tag{S13}$$

where

$$\mathbf{A}_q^{\text{2D}} = \left[ \mathbf{B}_a(x_{q0}, y_{q0}), \frac{\partial \mathbf{B}_a^o(x_{q0}, y_{q0})}{\partial x}, \frac{\partial \mathbf{B}_a^o(x_{q0}, y_{q0})}{\partial y} \right], \tag{S14a}$$

$$\boldsymbol{\zeta}_q^{\text{2D}} = \left[ s_q \boldsymbol{m}_q^T, s_q (\boldsymbol{m}_q^o)^T dx_q, s_q (\boldsymbol{m}_q^o)^T dy_q \right]^T, \tag{S14b}$$

$$\mathbf{B}_a = (\mathbf{B}(x_q, y_q, h_q = 0) + \mathbf{B}(x_q, y_q, h_q = 900\text{ nm}))/2, \tag{S14c}$$

and $\mathbf{B}_a$ is an average basis matrix combining the basis images at the top of the focal volume with the basis images at the bottom axial plane (Fig. S2). We use the RoSE-O algorithm [7], which is designed to detect and estimate the 2D locations and 3D orientations of SMs, to determine the number of emitters and an initial estimate of their 2D locations $(\hat{x}_q, \hat{y}_q)$.

In the second step, we obtain an initial estimate of the 3D orientations and 3D locations of individual emitters. For each detected emitter, we isolate an image of $21 \times 21$ pixels centered at the grid point $(\hat{x}_{q0}, \hat{y}_{q0})$ nearest to the estimated 2D location $(\hat{x}_q, \hat{y}_q)$ from the first step. Using each cropped image $\boldsymbol{I}_{\text{det}}$, algorithm 1 estimates the 15 parameters in $\boldsymbol{\zeta}_q$ simultaneously by minimizing the negative log-likelihood for $q^{\text{th}}$ emitter

$$\ell_{\text{NLL},q} = \sum_{i=1}^{N} \left[ \mathbf{A}_q \boldsymbol{\zeta}_q + \boldsymbol{b} - \boldsymbol{I}_{\text{det}} \odot \log\left( \mathbf{A}_q \boldsymbol{\zeta}_q + \boldsymbol{b} \right) \right]_i, \tag{S15}$$

where $[\cdot]_i$ represents the $i^{\text{th}}$ element of a vector. In step 3, the estimated $\boldsymbol{\zeta}_q$ for the Q SMs are combined as the initial value $\boldsymbol{\zeta}^0$. Algorithm 2, based on FISTA [8], is used to refine brightness, position, and orientation estimates $\boldsymbol{\zeta}_q$ simultaneously for all emitters based on the entire captured image by minimizing the negative log-likelihood for all Q SMs

$$\ell_{\text{NLL}} = \sum_i^N \left\{ \sum_q (\mathbf{A}_q \boldsymbol{\zeta}_q) + \boldsymbol{b} - \boldsymbol{I}_{\text{det}} \odot \log\left[ \sum_q (\mathbf{A}_q \boldsymbol{\zeta}_q) + \boldsymbol{b} \right] \right\}_i. \tag{S16}$$

**Algorithm 1** Estimating the 3D orientation and 3D location of individual emitters based on cropped images

**input:** $[\boldsymbol{I},\mathbf{A}_{\text{library}}]$
  ▷ $\mathbf{A}_{\text{library}} = [\mathbf{A}_1, ..., \mathbf{A}_n]$ is a set of basis matrices for emitters located at $[x, y, h] = \{[0, 0, \boldsymbol{h}_1],...,[0, 0, \boldsymbol{h}_n]\}$, where $\boldsymbol{h} = [-300,-250,...,1300]$ nm represents the locations of $n$ h-slices.
**for** $j \leftarrow 1$ **to** $n$ **do** ▷ $n$ is the number of h-slices.
  $\mathbf{A} \leftarrow \mathbf{A}_j$ ▷ $\mathbf{A}$ represents the current basis matrix.
  Crop $\mathbf{A}$ so that each basis image in $\mathbf{A}$ has a size of $21 \times 21$ pixels
  $\zeta^0_{q,j} \leftarrow \mathbf{A}^{-1}(\boldsymbol{I} - b)$
    ▷ Obtain an initial brightness, position, and orientation estimate for the $q^{\text{th}}$ emitter, assuming its image matches $\boldsymbol{A}_j$.
  Obtain a refined $\boldsymbol{\zeta}_{q,j}$, grid estimation $\boldsymbol{w}_{q,j}$, and loss value $\ell_{\text{NLL},q,j}$ from algorithm 2.
    ▷ Use $\zeta^0_{q,j}$ as the initial value, $[0, 0, h_j]$ as the initial grid location estimation $w^0_{q,j}$, and an iteration number of 30 for algorithm 2.
**end for**
**end for**
Obtain a final estimate $\boldsymbol{\zeta}_q$ and $\boldsymbol{w}_q$ from algorithm 2.
  ▷ Use $\boldsymbol{\zeta}_{q,j}$ and $w_{q,j}$ that has the smallest loss value $\ell_{\text{NLL},q,j}$ and an iteration number of 100 for algorithm 2.

**Algorithm 2** Estimating 3D orientation and 3D location using FISTA with backtracking

**input:** $[\boldsymbol{I},\boldsymbol{\zeta}^0,\boldsymbol{w}^0,\mathbf{A}_{\text{library}},\text{N}]$
Using $\mathbf{A}_{\text{library}}$, compute basis matrix $\mathbf{A}$ centered at the estimated initial grid location $\boldsymbol{w}^0$.
$\boldsymbol{w}_0 \leftarrow \boldsymbol{w}^0$, $\boldsymbol{\zeta}_0 \leftarrow \boldsymbol{\zeta}^0$, $\boldsymbol{v}_1 \leftarrow \boldsymbol{\zeta}^0$ ▷ $\boldsymbol{v}_1$ is a nearby point of $\boldsymbol{\zeta}^0$ that will be used to accelerate the optimization.
$t_1 \leftarrow 1$, and take $L_0 > 0$ and $\eta > 1$.
  ▷ $L_0$ denotes a constant chosen for initializing the step size, and $\eta$ is a constant chosen to adjust step sizes in future steps.
**for** $k \leftarrow 1$ **to** $N$ **do** ▷ N is the total number of iterations
  Calculate the best step size by finding the smallest integer $i_k > 0$ such that with $L_k = \eta^{i_k} L_{k-1}$: $\ell_{\text{NLL}}(\boldsymbol{v}_{k,L_k}) \leq \ell_{\text{NLL}}(\boldsymbol{v}_k) + \nabla_{\boldsymbol{v}}\ell_{\text{NLL}}(\boldsymbol{v}_k)^T [\boldsymbol{v}_{k,L_k} - \boldsymbol{v}_k] + \frac{L_k}{2}\|\boldsymbol{v}_{k,L_k} - \boldsymbol{v}_k\|_2^2$
    ▷ $\nabla_{\boldsymbol{v}}\ell_{\text{NLL}}(\boldsymbol{v}_k) \leftarrow \mathbf{A}^T(1 - (\boldsymbol{I} \oslash (\mathbf{A}\boldsymbol{v}_k + \boldsymbol{b})))$, and operator $\oslash$ represents element-wise division of two vectors.
  $\boldsymbol{v}_{k,L_k} = P_{L2}\left(P_{L1}\left(\boldsymbol{v}_k - \frac{1}{L_k}\nabla_{\boldsymbol{v}}\ell_{\text{NLL}}(\boldsymbol{v}_k)\right)\right)$, and $P_{L1}$ and $P_{L2}$ are projection operators shown in Eqn. S20 and Eqn. S21
  $L_k \leftarrow \eta^{i_k} L_{k-1}$
  $\boldsymbol{\zeta}_k \leftarrow P_{L2}(P_{L1}\left(\boldsymbol{v}_k - \frac{1}{L_k}\nabla_v\ell_{\text{NLL}}(\boldsymbol{v}_k)\right))$
  Based on $\boldsymbol{\zeta}_k$ and $\boldsymbol{w}_{k-1}$, compute $[\hat{x}, \hat{y}, \hat{h}]$ and its closest grid point $\boldsymbol{w}_k = [\hat{x}_0, \hat{y}_0, \hat{h}_0]$ using Eqn. S17
  Compute $\mathbf{A}$ centered at the new grid location $\boldsymbol{w}_k$ using $\mathbf{A}_{\text{library}}$.
  Compute $\boldsymbol{\zeta}_{k-1,\text{new}}$ and $\boldsymbol{\zeta}_{k,\text{new}}$ based on the new grid location $\boldsymbol{w}_k$.
    ▷ $\boldsymbol{\zeta}_{k-1,\text{new}} = \boldsymbol{\zeta}_{k-1} + [\boldsymbol{o}_0, o_1\boldsymbol{\zeta}_{k-1,[1:3]}, o_2\boldsymbol{\zeta}_{k-1,[1:3]}, o_3\boldsymbol{\zeta}_{k-1,[1:3]}]$, $\boldsymbol{\zeta}_{k,\text{new}} = \boldsymbol{\zeta}_k + [\boldsymbol{o}_0, o_1\boldsymbol{\zeta}_{k,[1:3]}, o_2\boldsymbol{\zeta}_{k,[1:3]}, o_3\boldsymbol{\zeta}_{k,[1:3]}]$, where $\boldsymbol{o}_0 = [0, 0, 0, 0, 0, 0]$, $o_1 = (\boldsymbol{w}_{k-1,1} - \boldsymbol{w}_{k,1})$, $o_2 = (\boldsymbol{w}_{k-1,2} - \boldsymbol{w}_{k,2})$, $o_3 = (\boldsymbol{w}_{k-1,3} - \boldsymbol{w}_{k,3})$
  $\boldsymbol{\zeta}_{k-1} \leftarrow \boldsymbol{\zeta}_{k-1,\text{new}}$, and $\boldsymbol{\zeta}_k \leftarrow \boldsymbol{\zeta}_{k,\text{new}}$
  $t_k \leftarrow \frac{1+\sqrt{1+4t_k^2}}{2}$
  $v_{k+1} \leftarrow \boldsymbol{\zeta}_k + \frac{t_k - 1}{t_{k+1}}(\boldsymbol{\zeta}_k - \boldsymbol{\zeta}_{k-1})$
**end for**
**end for**

The 3D locations $(\hat{x}_q, \hat{y}_q, \hat{h}_q)$ and orientational moments $\hat{\boldsymbol{m}}_q$ of molecule $q$ can be extracted from $\boldsymbol{\zeta}_q$ using

$$\hat{s}_q = \sum_{i=1}^{3} \zeta_{q,i}, \tag{S17a}$$

$$\hat{\boldsymbol{m}}_q = \frac{1}{\hat{s}_q}\boldsymbol{\zeta}_{q,1:6}, \tag{S17b}$$

$$\hat{x}_q = \hat{x}_{q0} + \frac{\sum_{i=7}^{9} \zeta_{q,i}}{\sum_{i=1}^{3} \zeta_{q,i}}, \tag{S17c}$$

$$\hat{y}_q = \hat{y}_{q0} + \frac{\sum_{i=10}^{12} \zeta_{q,i}}{\sum_{i=1}^{3} \zeta_{q,i}}, \tag{S17d}$$

$$\hat{h}_q = \hat{h}_{q0} + \frac{\sum_{i=13}^{15} \zeta_{q,i}}{\sum_{i=1}^{3} \zeta_{q,i}}, \tag{S17e}$$

where $\zeta_{q,i}$ represents the $i^{\text{th}}$ element of vector $\boldsymbol{\zeta}_q$. The estimated second-moment vectors $\hat{\boldsymbol{m}}_q$ are next projected to first-moment orientation space ($\hat{\phi}$, $\hat{\theta}$, $\hat{\Omega}$) using a weighted least-square estimator as follows:

$$\left(\hat{\phi}, \hat{\theta}, \hat{\Omega}\right) = \underset{\phi',\theta',\Omega'}{\arg\min}\left(\hat{\boldsymbol{m}}_q - \boldsymbol{m}_q(\phi', \theta', \Omega')\right)^T \mathbf{K}^{-1} \left(\hat{\boldsymbol{m}}_q - \boldsymbol{m}_q(\phi', \theta', \Omega')\right). \tag{S18}$$

Note that weighting by a Fisher information matrix ($\mathbf{K}^{-1}$) ensures that more weight is given to the second moments $m_i$ for which pixOL demonstrates superior precision.

When estimating $\boldsymbol{\zeta}_q$, we use two sequential projection operations to enhance the convexity and guarantee that the second-moment $\hat{\boldsymbol{m}}_q$ estimates are physically meaningful. The first projection $P_{L1}$ enforces the first three brightness-weighted moments $[\zeta_{q,1}, \zeta_{q,2}, \zeta_{q,3}]$ to be positive. It also ensures that the off-grid distances $[dx_q, dy_q, dh_q]$ are smaller than a threshold $t$, since a large off-grid distance will reduce the robustness of the first-order polynomial approximation (Eqn. S10) [9]. Parameters $[\zeta_{q,i}, d\boldsymbol{w}_q^i]$ related to each orientational moment $m_{q,i}$ are projected separately, where

$$i \in \{1,2,3\}, \tag{S19a}$$
$$d\boldsymbol{w}_q^1 = \boldsymbol{\zeta}_{q,[7,10,13]}, \tag{S19b}$$
$$d\boldsymbol{w}_q^2 = \boldsymbol{\zeta}_{q,[8,11,14]}, \tag{S19c}$$
$$d\boldsymbol{w}_q^3 = \boldsymbol{\zeta}_{q,[9,12,15]}. \tag{S19d}$$

We write this projection as

$$P_{L1}([\zeta_{q,i}, d\boldsymbol{w}_q^i]) = \begin{cases} 0, & \text{if } \|d\boldsymbol{w}_q^i\|_2 \le \frac{-\zeta_{q,i}}{t} \\ (\zeta_{q,i}, d\boldsymbol{w}_q^i), & \text{if } \|d\boldsymbol{w}_q^i\|_2 \le \zeta_{q,i} t \\ (\frac{\zeta_{q,i}+t\|d\boldsymbol{w}_q^i\|_2}{1+t^2}, & \\ \quad \frac{\zeta_{q,i}+t\|d\boldsymbol{w}_q^i\|_2}{1+t^2} \frac{t d\boldsymbol{w}_q^i}{\|d\boldsymbol{w}_q^i\|_2}), & \text{if } \|d\boldsymbol{w}_q^i\|_2 \ge \zeta_{q,i} t, \end{cases} \tag{S20}$$

and $t = 117$ nm in our case.

The second projection $P_{L2}$ ensures that the second-moment $\hat{\boldsymbol{m}}_q$ estimates correspond to a wobble $\Omega$ that is physically meaningful, given by

$$P_{L2}(\boldsymbol{\zeta}_q) = [\boldsymbol{\zeta}_{q,1:3}, k\boldsymbol{\zeta}_{q,4:6}, \boldsymbol{\zeta}_{q,7:15}], \tag{S21}$$

where

$$k = \frac{\max(0, \min(1.5e - 0.5, 1))}{1.5e - 0.5}, \tag{S22}$$

and $e$ is the largest eigenvalue of the second moment matrix.

### III. IMAGING SYSTEM CALIBRATION

We calibrate the imaging model in Section II to the DSF of our imaging system by using fluorescent beads (100-nm diameter red 580/605 FluoSpheres, Invitrogen F8801). Images are captured by scanning the objective's nominal focal plane from $z = -790$ nm to $z = 610$ nm with a step size of 50 nm (Fig. S7(e), Fig. S8(f), Fig. 2(e)). At each plane, 11 images are taken.

A phase-retrieval algorithm [10] is used to retrieve the experimental phase mask, calibrated pixOL $\tilde{\boldsymbol{P}}$. To accurately characterize the optical aberrations of our microscope, the phase masks of the two polarized channels are estimated independently (Fig. S7(b,c), Fig. S8(c,d)). The simulated DSFs using the calibrated phase masks $\tilde{\boldsymbol{P}}$ match the experimental DSFs very well (Fig. S7(f), Fig. S8(g)).

We performed phase retrieval for both the pixOL phase mask and its conjugate mask (pixOL*). We notice that the pixOL phase mask's DSF has a large aberration when the objective is focused below the coverslip ($z > 0$, Fig. S7(e)). While aberrations also exist in the pixOL* experimental DSF at the same locations, its mismatch relative to the ideal pixOL DSF is more modest and is similar to spherical aberration (Fig. S8(f)).

### IV. QUANTIFYING THE PRECISION OF MEASURING 3D ORIENTATION AND 3D POSITION

The Fisher information matrix quantifies the amount of information contained within any DSF regarding the parameters we want to estimate. The shape of the pixOL DSF contains information of both the emitter's axial location and its 3D orientation. Due to the discrete sampling and finite size of the camera pixels, the off-grid distances $[dx, dy, dh]$ related to the emitter's location $[x, y, h]$ relative to the closest grid point $[x_0, y_0, h_0]$ also affect the shape of the DSF image. Therefore, the covariance between any pair of measurement parameters $[x, y, h, \theta, \phi, \Omega]$ representing position and orientation is nonzero. Thus, to properly quantify the best-possible precision of measuring 3D position and 3D orientation, while also considering correlations thereof, we calculate Fisher information matrices for joint estimation of 3D position and 3D orientation (Fig. 2(a-d)).

For Fig. 2(a,b), we calculate a $6 \times 6$ Fisher information matrix that includes 3 location and 3 orientation parameters. By inverting the Fisher information matrix, we obtain the CRB matrix $\mathbf{K}$, whose diagonal elements quantify the estimation variance of 3D orientation $[\theta, \phi, \Omega]$ and 3D location $[x, y, h]$. To quantify the precision of estimating the mean orientation $[\theta, \phi]$, we calculate the angular precision $\sigma_\delta$, which is the half-angle of the uncertainty cone for estimating the mean orientation direction [11]. It is a summary metric that combines the precision $\sigma_\theta$ of measuring $\theta$ and the standard deviation $\sigma_\phi$ of measuring $\phi$, given by

$$\sigma_\delta = 2 \arcsin \left( \sqrt{\frac{\sin(\theta)\sqrt{\det(\mathbf{K}_{4:5,4:5})}}{4\pi}} \right), \tag{S23}$$

where $\mathbf{K}_{4:5,4:5}$ is a $2 \times 2$ sub-matrix of $\mathbf{K}$ representing 3D orientation $[\theta, \phi]$ precision formed by fourth and fifth rows and columns. We sample orientation space $(\theta, \phi)$ uniformly using

$$\phi = 2\pi v, \tag{S24}$$
$$\theta = \arccos(2u - 1), \tag{S25}$$

where $v, u$ are uniformly distributed on (0,1). The angular precision $\sigma_\delta$ and wobbling angle precision $\sigma_\Omega$ is then averaged over all $\theta$ in Fig. 2(a,b).

To quantify 3D location precision for isotropic emitters (Fig. 2(c,d)), we build a $9 \times 9$ Fisher information matrix, which includes the 3 location parameters and 6 orientational moments. This formulation avoids the undefined mean orientation direction $(\theta, \phi)$ of these emitters.

The correlation between the 3D orientation and 3D location will degrade estimation precision in general. We quantify the difference ratio $Y$ between the precision calculated with orientation-position correlation and without using

$$Y(\sigma) = \frac{\sigma_{\text{with correlation}} - \sigma_{\text{without correlation}}}{\sigma_{\text{without correlation}}}, \tag{S26}$$

where $\sigma_{\text{without correlation}}$ is calculated using matrices $\mathbf{K}$ that quantify solely 3D localization precision or 3D orientation measurement precision. We noticed that the performance of large DSFs (i.e., those that are greater than 3 times the size of the standard PSF) is less influenced by orientation-position correlations (Fig. S4).

### V. QUANTIFYING ESTIMATION BIAS WHEN MEASURING 3D ORIENTATION AND 3D POSITION

To determine if correlations between measurements of 3D orientation and 3D position influence estimation performance, we simulated images of emitters with various orientations and axial locations and a nominal focal plane set to $z = -580$ nm (Figs. S9, S10, and S11). For each configuration, 200 independent images are simulated using the forward model (Eqn. 1) and estimated using algorithm described in Section II. Overall, pixOL shows excellent precision for measuring 3D orientation and 3D position. However, in some regions of the orientation domain, we notice low estimation precision and accuracy for both 3D orientation and 3D position. Plotting the estimates in one of these regions, we notice strong correlations between biases in 3D orientations and biases in 3D locations (Fig. S13(a,b)). For example, simulated noiseless images of an emitter at the ground truth position $(x_0, y_0, h_0) = (0, 0, 700)$ nm and orientation $(\theta_0, \phi_0, \Omega_0) = (38^\circ, 162^\circ, 0)$ are very similar to those of a biased estimate of position and orientation (Fig. S13(c,d)). However, differences between the noisy images are still discernible. We suspect that position-orientation correlations create local minima in the likelihood space, making robust estimation difficult. We believe that a more powerful algorithm that is able to explore efficiently 6D position-orientation space to find the global minimum negative log-likelihood will increase estimation precision and remove bias (Fig. S13(e)).

### VI. PREPARING 3D SPHERICAL SUPPORTED LIPID BILAYERS

Supported lipid bilayers (SLBs) were formed by fusing vesicles on silica beads. A lipid mixture containing 0.286 mg/mL DPPC (di(16:0) phosphatidylcholine, Avanti Polar Lipids 850355) and 0.1 mg/mL cholesterol (Sigma Aldrich C8667) within Tris buffer (100 mM NaCl, 3 mM $Ca^{2+}$, 10 mM Tris, pH 7.4) is first incubated in a water bath at a temperature of 65 °C. Simultaneously, 1 mg/mL silica beads (2.0 $\mu$m diameter, Bangs Laboratories SS04002) are also incubated in the water bath for ~10 minutes. After two solutions reach to thermal equilibrium, 30 $\mu$L of the lipid mixture and 90 $\mu$L of the silica beads are mixed together and stay in the water bath for another 30 minutes. During the 30 minutes, the mixture is vortexed every 5 minutes. The mixture is then moved outside the water bath together with around 200 mL of warm water to allow it to cool down to room temperature for about 1 hour. During this cooling process, the mixture is vortexed every 5 minutes to allow the lipid bilayer to coat the bead uniformly.

After the mixture cool down to the room temperature, we remove the excess vesicles using six successive 5-min centrifuge spins at 500 RPM. The supernatant is discarded and replaced with imaging buffer (100 mM NaCl, 10 mM Tris, pH 7.4) after each spin.

### VII. IMAGING SPHERICAL LIPID BILAYERS USING NILE RED

Nile red (32 nM) is added to the imaging buffer to probe the morphology and composition of the spherical lipid bilayers. We use circularly polarized illumination (1533 W/cm$^2$ or 30 mW at 561 nm at the sample) to excite NR. Fluorescence is collected using an objective lens (OLYMPUS UPLSAPO100XOPSF, NA 1.4) and passes through a dichroic mirror (Di01-R488/561, Semrock) and bandpass filter (FF01-523/610, Semrock).

We captured 20000 frames with an exposure time of 110 ms. To compensate for axial drift of the microscope during long-term imaging, the objective is refocused every 2000 frames. We use a nominal focal plane (NFP) of $z = -350$ nm for the DPPC+chol bead and an NFP of $z = -500$ nm for the DPPC-only bead. The lateral drift of the microscope is corrected after imaging. We aligned the lateral centers of each sphere using SM position estimates averaged in groups of 2000 frames. With these experimental conditions, we observe single-molecule blinking with properties shown in Fig. S15.

To quantify the localization precision for NRs, we plot the NR lateral positions $r$ relative to the sphere's center within each $h$ slice (Figs. S16(d) and S17(d)). We compare the experimental distribution to a theoretical distribution $p_{\text{theo}}$ calculated by convolving the lateral positions $p_{\text{sphere}}$ representing the spherical surface with the expected lateral localization distribution $p_{\text{pixOL}^*}$ of pixOL* based on the CRB when normal focaol plane equals to -580 nm, i.e.,

$$p_{\text{theo}}(r) = p_{\text{sphere}} \circledast p_{\text{pixOL}^*}, \tag{S27}$$

where

$$p_{\text{sphere}}(r) = \frac{R}{(\sqrt{R^2 - r^2})}, \tag{S28a}$$

$$p_{\text{pixOL}^*}(r) = \frac{1}{\sigma_L(\sqrt{R^2 - r^2})\sqrt{2\pi}} \times \exp\left[-\frac{1}{2}\left(\frac{r}{\sigma_L(\sqrt{R^2 - r^2})}\right)^2\right], \tag{S28b}$$

$R$ is the radius of the sphere, $\sigma_L(\sqrt{R^2 - r^2})$ is the lateral precision of pixOL* for an emitter on the sphere's surface with $h = R - \sqrt{R^2 - r^2}$, and $\circledast$ represents convolution operation.

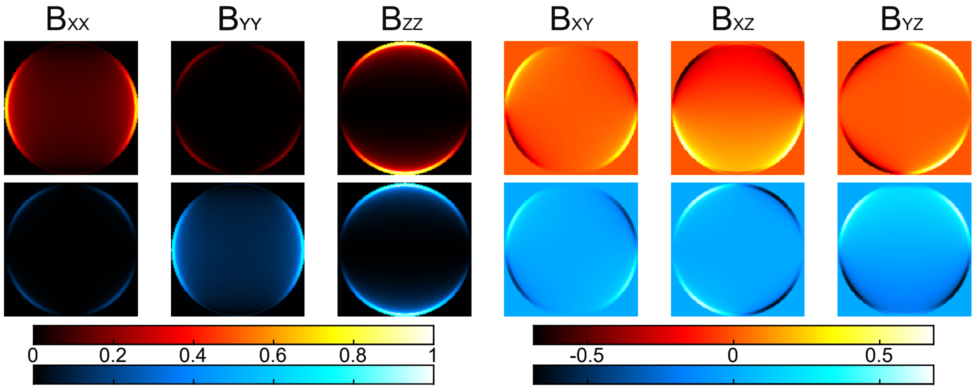

FIG. S1. Basis images $\mathbf{B}_{\text{BFP}}$ at the back-focal plane (BFP) of an x-y polarized microscope (red: x-polarized, blue: y-polarized, Fig. 1(a)). The radius of the circular support in the BFP corresponds to the finite numerical aperture of the microscope's objective lens. The image intensities are normalized relative to the brightest basis image ($\mathbf{B}_{\text{XX}}$). Colorbars: normalized intensity.

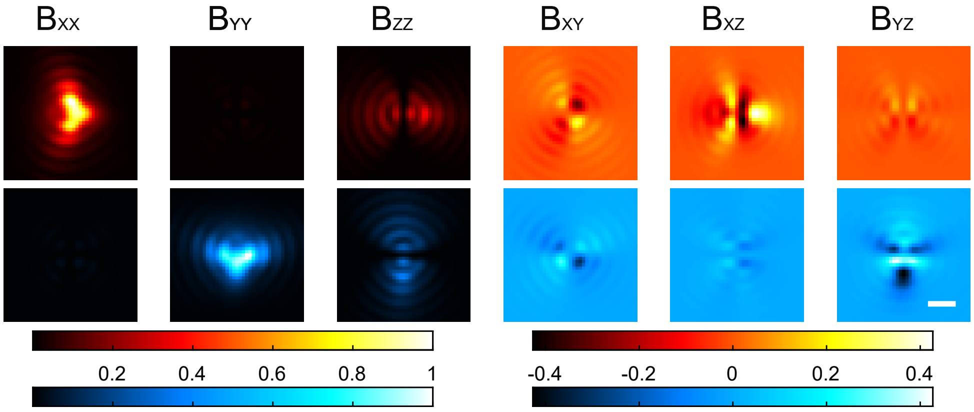

FIG. S2. Average basis images $\mathbf{B}_a$ (red: x-polarized, blue: y-polarized) used for detecting single molecules (Eqn. S14c). Basis images for an emitter at $h = 0$ nm and $h = 900$ nm are averaged together. The image intensities are normalized relative to the brightest basis image ($\mathbf{B}_{\text{XX}}$). Colorbars: normalized intensity. Scalebar: 500 nm.

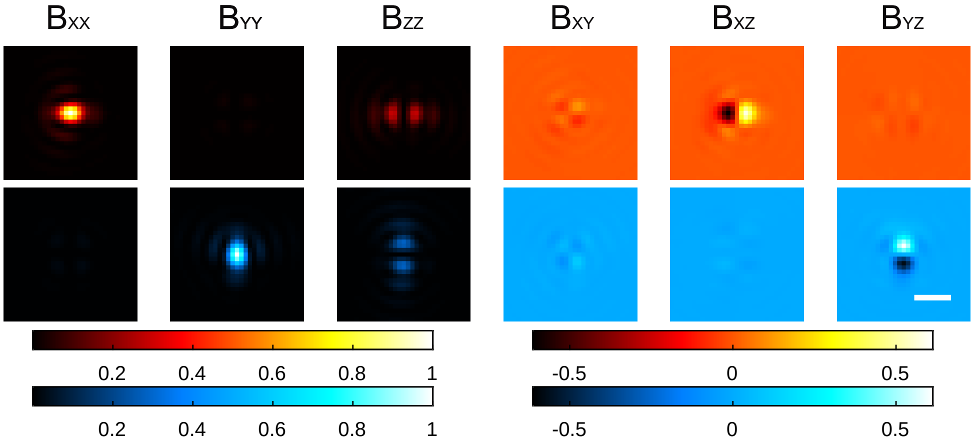

FIG. S3. Image plane basis images $\mathbf{B}$ corresponding to the polarization-sensitive imaging system (red: x-polarized, blue: y-polarized) and pixOL phase mask shown in Fig. 1(a,c) for an in-focus emitter. The image intensities are normalized relative to the brightest basis image ($\mathbf{B}_{\text{XX}}$). Colorbars: normalized intensity. Scalebar: 500 nm.

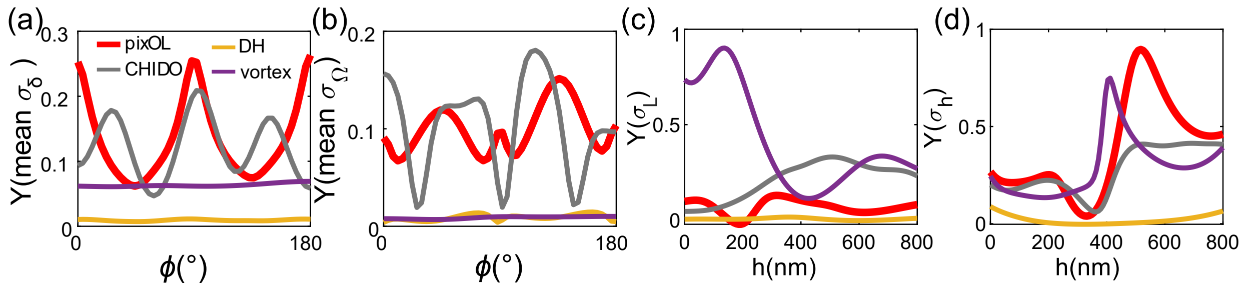

FIG. S4. Influence of 3D orientation-3D position correlations on measurement precision. (a) Difference ratio $Y$, which quantifies the difference in measurement precision accounting for orientation-position correlations vs. ignoring them (Eqn. S26), for orientation precision $\sigma_\delta$, averaged over all polar orientations $\theta$. (b) Difference ratio $Y$ for wobble angle precision $\sigma_\Omega$. (c) Difference ratio $Y$ for lateral localization precision $\sigma_L$. (d) Difference ratio $Y$ for axial localization precision $\sigma_h$. In-focus SMs with fixed orientation ($\Omega = 0$ sr) are used for (a,b); isotropic emitters ($\Omega = 2\pi$ sr) are used for (c,d). Emitters are immersed in water (1.33 refractive index) with 2500 signal photons and 3 background photons per pixel.

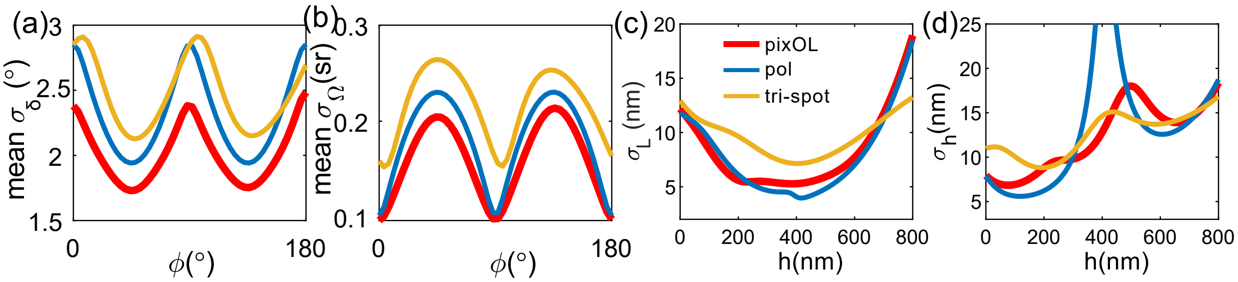

FIG. S5. Estimation precision of pixOL compared to techniques designed solely for 3D orientation measurements. Red: pixOL, blue: x-y polarized standard PSF (pol) defocused at 200 nm above the coverslip [12], yellow: tri-spot [13]. (a) Mean angular standard deviation $\sigma_\delta$ (MASD) averaged uniformly over all $\theta$. MASD quantifies the combined precision of measuring $\theta$ and $\phi$ as the half-angle of a cone representing orientation uncertainty (Eqn. S23) [11]. (b) Mean wobble angle precision $\sigma_\Omega$ averaged uniformly over all $\theta$. (c,d) Localization precisions $\sigma_L$ and $\sigma_h$ for measuring (c) lateral position $l$ and (d) axial location $h$ above the interface, respectively. MASD and $\sigma_\Omega$ are calculated for in-focus SMs with fixed orientation ($\Omega = 0$ sr); localization precisions are for isotropic emitters ($\Omega = 2\pi$ sr). Emitters are immersed in water (1.33 refractive index) with 2500 signal photons and 3 background photons per pixel.

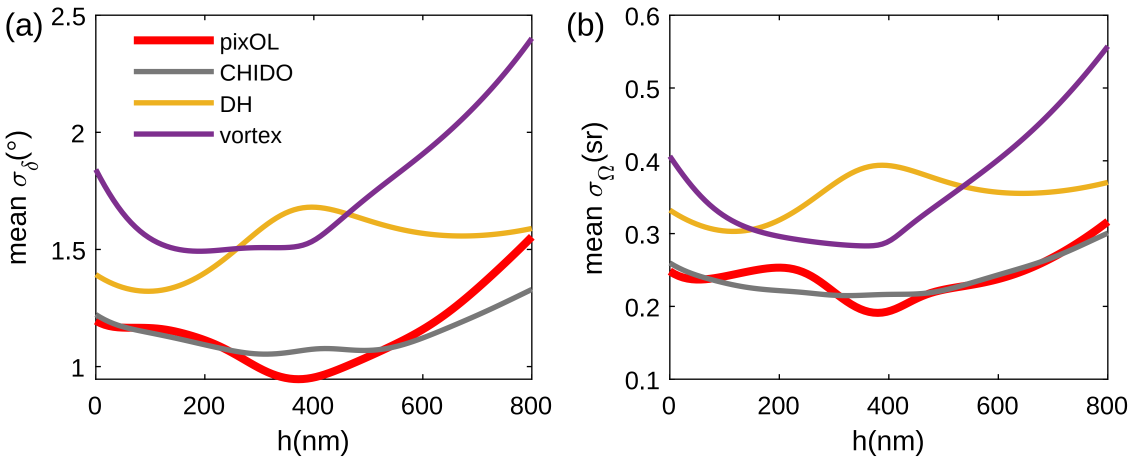

FIG. S6. Orientation estimation precision of pixOL for emitters at various axial locations $h$ compared to techniques designed for 3D orientation and 3D position measurements. Red: pixOL, grey: CHIDO [14], yellow: double helix (DH) [15], purple: unpolarized vortex [16]. (a) Mean angular standard deviation $\sigma_\delta$ (MASD) averaged uniformly over all $\theta$ and $\phi$. MASD quantifies the combined precision of measuring $\theta$ and $\phi$ as the half-angle of a cone representing orientation uncertainty (Eqn. S23) [11]. (b) Mean wobble angle precision $\sigma_\Omega$ averaged uniformly over all $\theta$ and $\phi$. Emitters have fixed orientations ($\Omega = 0$ sr), and are immersed in water (1.33 refractive index) with 2500 signal photons and 3 background photons per pixel.

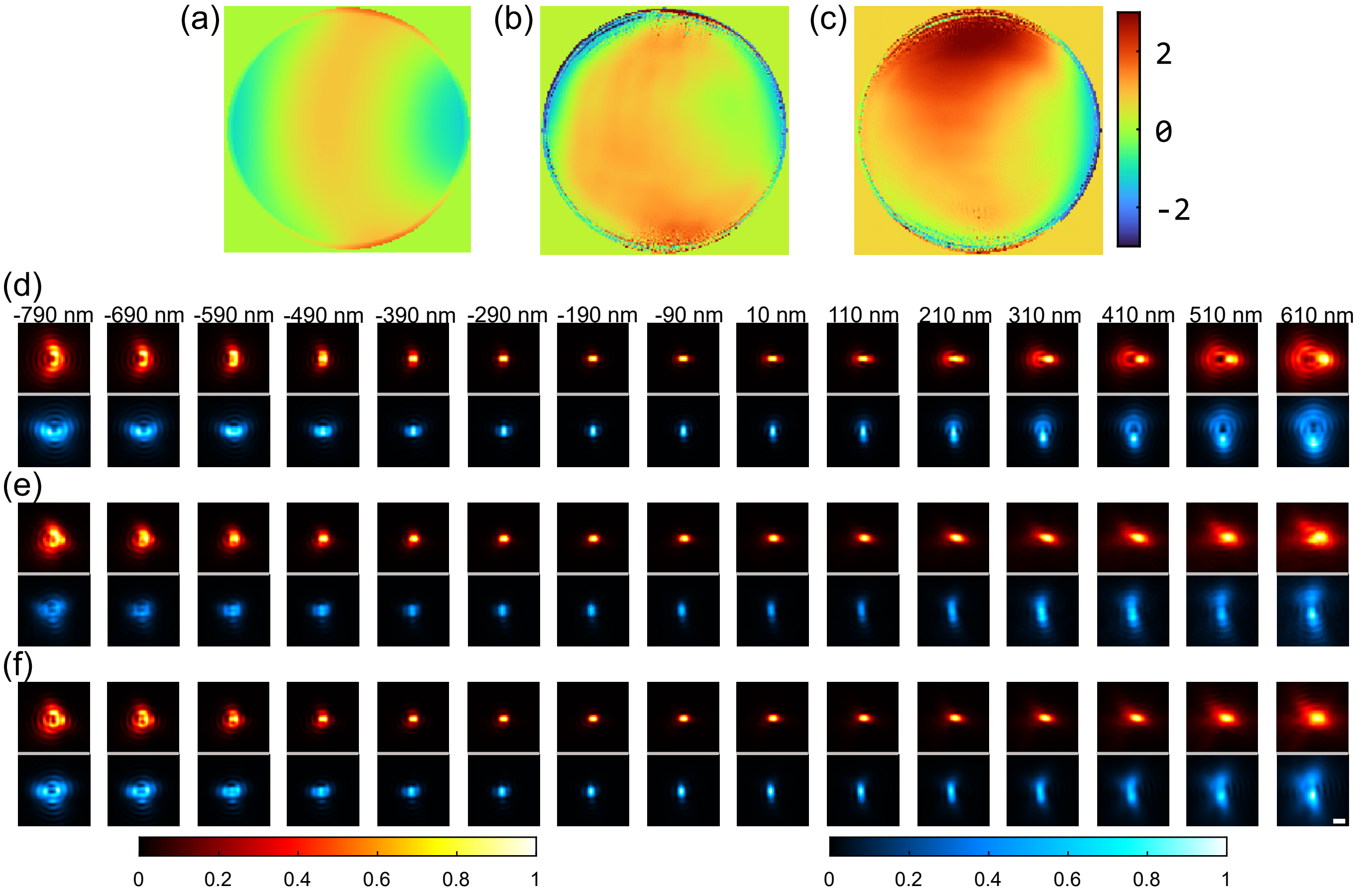

FIG. S7. Calibration of the pixOL phase mask. (a) Perfect pixOL phase mask. (b,c) Calibrated experimental pupil phase patterns from the (b) x and (c) y polarization channels. Colorbar: phase (rad). (d) Ideal pixOL DSFs (red: x-polarized, blue: y-polarized) for various defocus positions $z = -790$ nm to $z = 610$ nm. (e) Experimental DSFs. (f) Simulated DSFs using the calibrated phase masks in (b) and (c). The intensities of each red-blue image pair are normalized. Colorbar: normalized intensity. Scalebar: 400 nm.

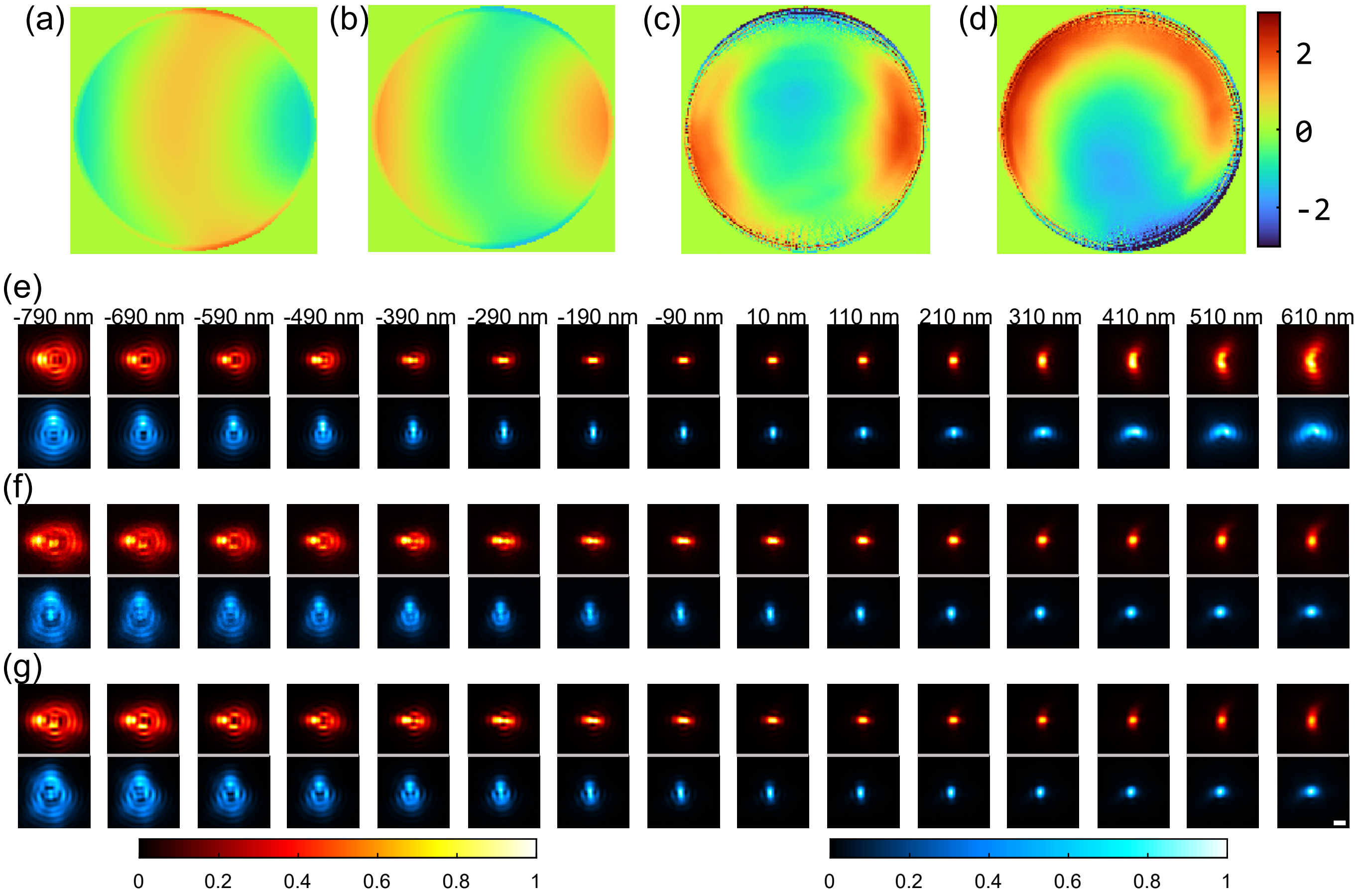

FIG. S8. Calibration of the conjugate pixOL phase mask (pixOL*). (a) Perfect pixOL phase mask. (b) Perfect pixOL* phase mask. (c,d) Calibrated experimental pupil phase patterns from the (c) x and (d) y polarization channels. Colorbar: phase (rad). (e) Ideal pixOL* DSFs (red: x-polarized, blue: y-polarized) for various defocus positions $z = -790$ nm to $z = 610$ nm. (f) Experimental DSFs. (g) Simulated DSFs using the calibrated phase masks in (c) and (d). The intensities of each red-blue image pair are normalized. Colorbar: normalized intensity. Scalebar: 400 nm.

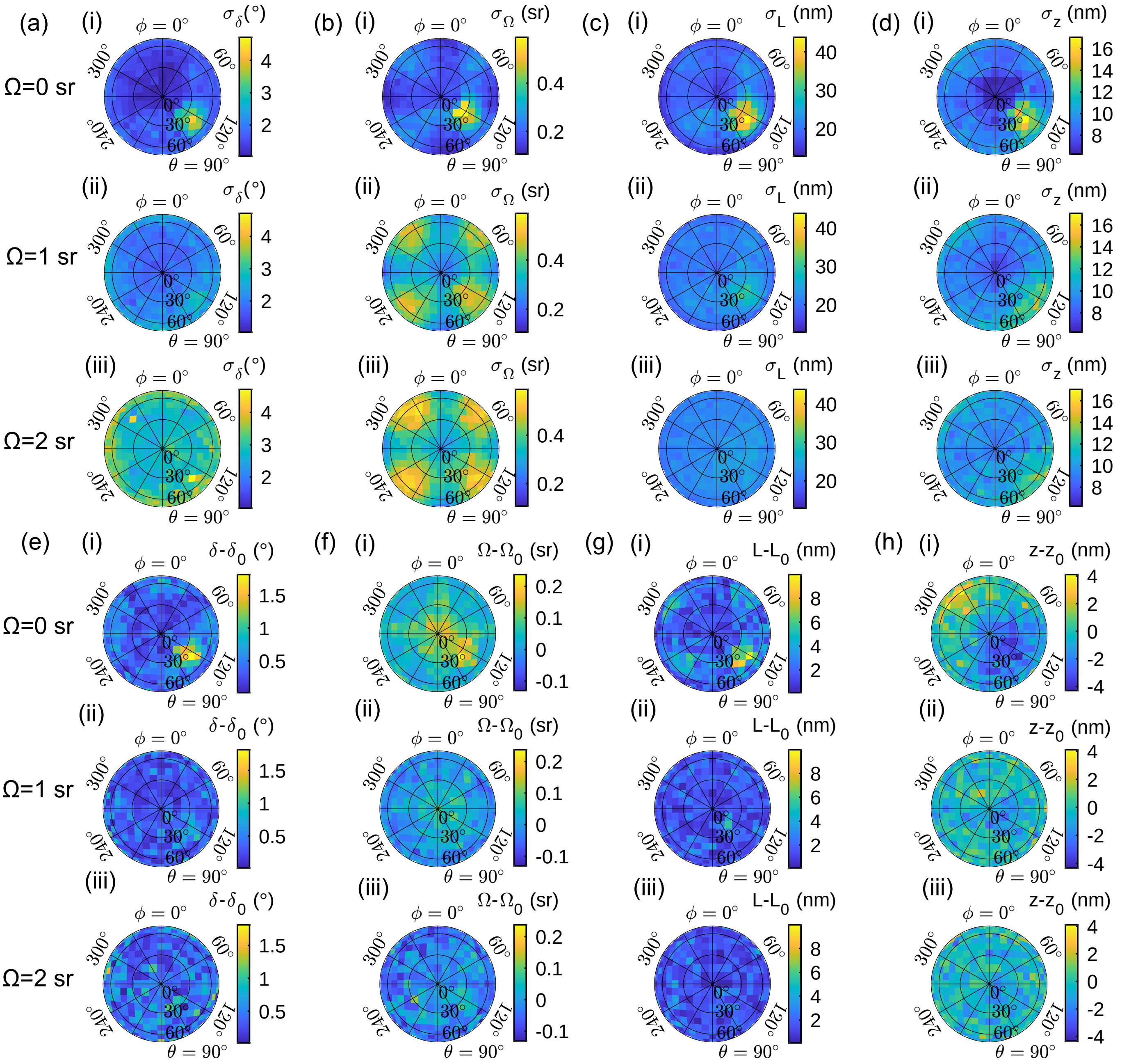

FIG. S9. Precision and accuracy of estimating 3D orientation and 3D position for emitters at $h = 0$ nm with a nominal focal plane placed at $z = -580$ nm. (a-d) Precision of measuring 3D orientation (a) $\sigma_\delta$ (deg), (b) $\sigma_\Omega$ (sr), and 3D location (c) $\sigma_L$ (nm), (d) $\sigma_z$ (nm) at various orientations ($\Omega \in \{0, 1, 2\}$ sr, $\theta \in [0°, 90°]$, $\phi \in [0°, 360°)$). (e-h) Accuracy of measuring 3D orientation (e) $\delta - \delta_0$ (deg), (f) $\Omega - \Omega_0$ (sr), and 3D location (g) $L - L_0$ (nm), (h) $z - z_0$ (nm). At each orientation, 200 independent images were generated for emitter within water located at $h = 0$ nm with 2500 signal photons in total and 3 background photons per pixel detected. The 3D orientation and 3D position are estimated as described in Section II.

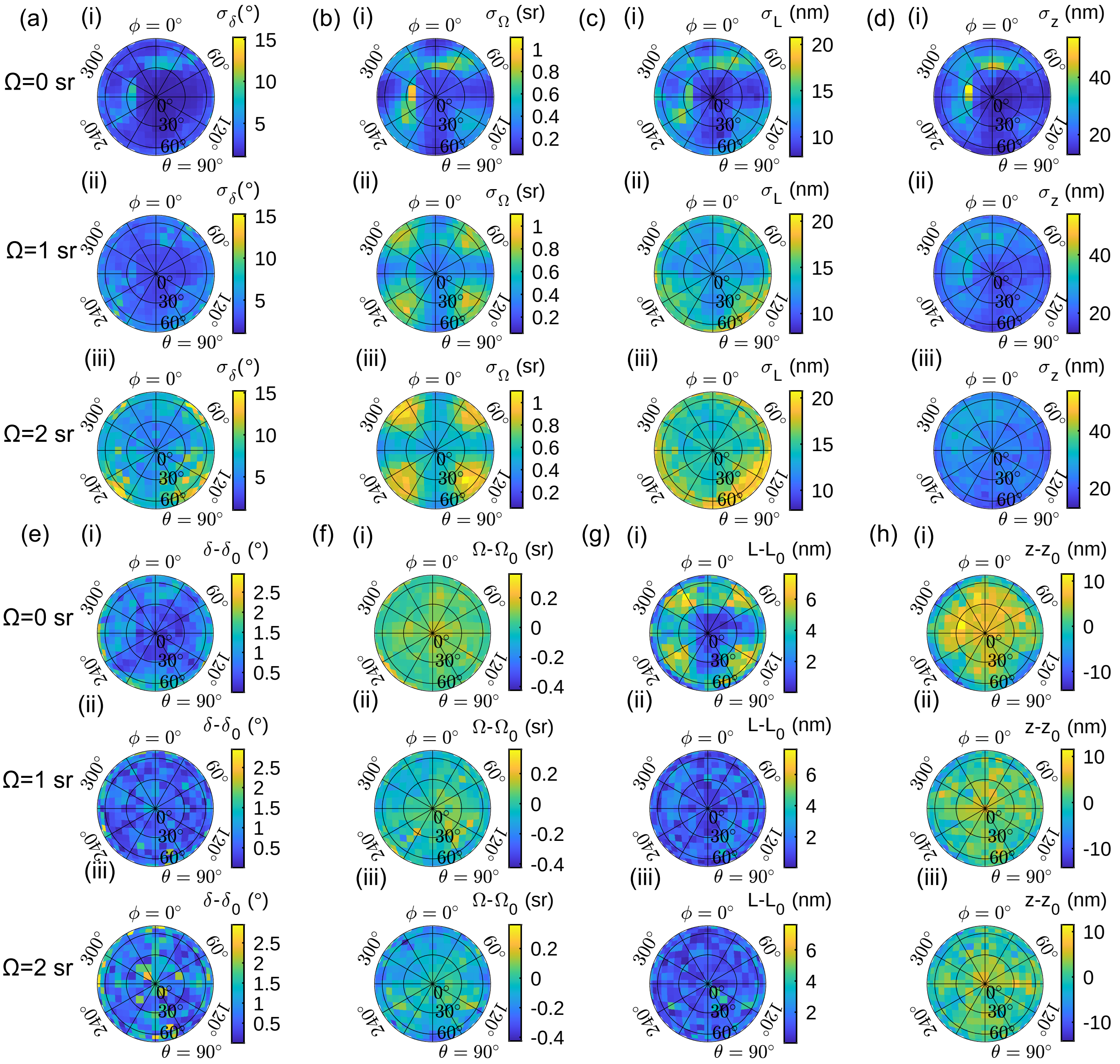

FIG. S10. Precision and accuracy of estimating 3D orientation and 3D position for emitters at $h = 400$ nm with a nominal focal plane placed at $z = -580$ nm. (a-d) Precision of measuring 3D orientation (a) $\sigma_\delta$ (deg), (b) $\sigma_\Omega$ (sr), and 3D location (c) $\sigma_L$ (nm), (d) $\sigma_z$ (nm) at various orientations ($\Omega \in \{0, 1, 2\}$ sr, $\theta \in [0^\circ, 90^\circ]$, $\phi \in [0^\circ, 360^\circ)$). (e-h) Accuracy of measuring 3D orientation (e) $\delta - \delta_0$ (deg), (f) $\Omega - \Omega_0$ (sr), and 3D location (g) $L - L_0$ (nm), (h) $z - z_0$ (nm). At each orientation, 200 independent images were generated for emitter within water located at $h = 400$ nm with 2500 signal photons in total and 3 background photons per pixel detected. The 3D orientation and 3D position are estimated as described in Section II.

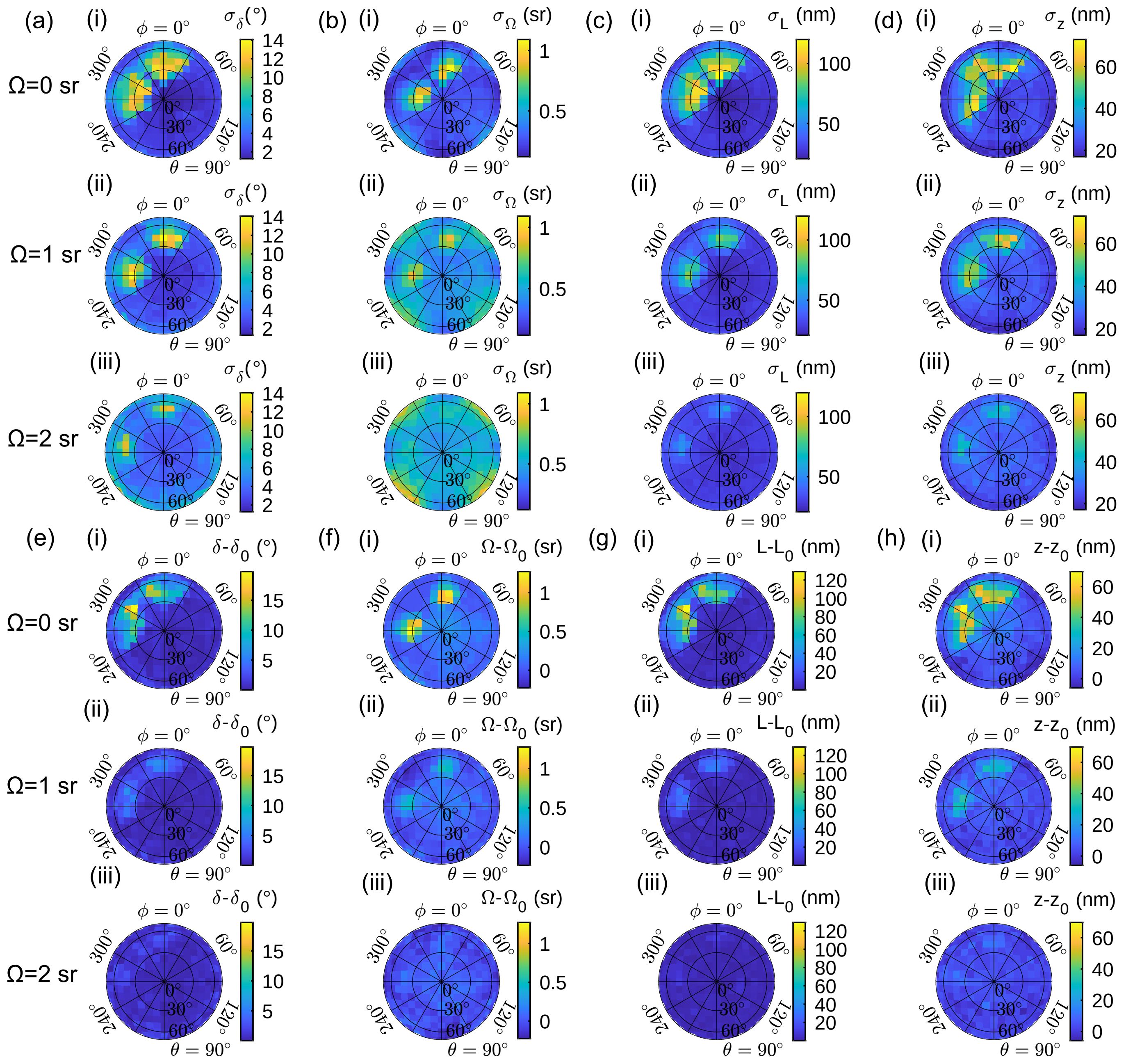

FIG. S11. Precision and accuracy of estimating 3D orientation and 3D position for emitters at $h = 700$ nm with a nominal focal plane placed at $z = -580$ nm. (a-d) Precision of measuring 3D orientation (a) $\sigma_\delta$ (deg), (b) $\sigma_\Omega$ (sr), and 3D location (c) $\sigma_L$ (nm), (d) $\sigma_z$ (nm) at various orientations ($\Omega \in \{0, 1, 2\}$ sr, $\theta \in [0°, 90°]$, $\phi \in [0°, 360°)$). (e-h) Accuracy of measuring 3D orientation (e) $\delta - \delta_0$ (deg), (f) $\Omega - \Omega_0$ (sr), and 3D location (g) $L - L_0$ (nm), (h) $z - z_0$ (nm). At each orientation, 200 independent images were generated for emitter within water located at $h = 700$ nm with 2500 signal photons in total and 3 background photons per pixel detected. The 3D orientation and 3D position are estimated as described in Section II.

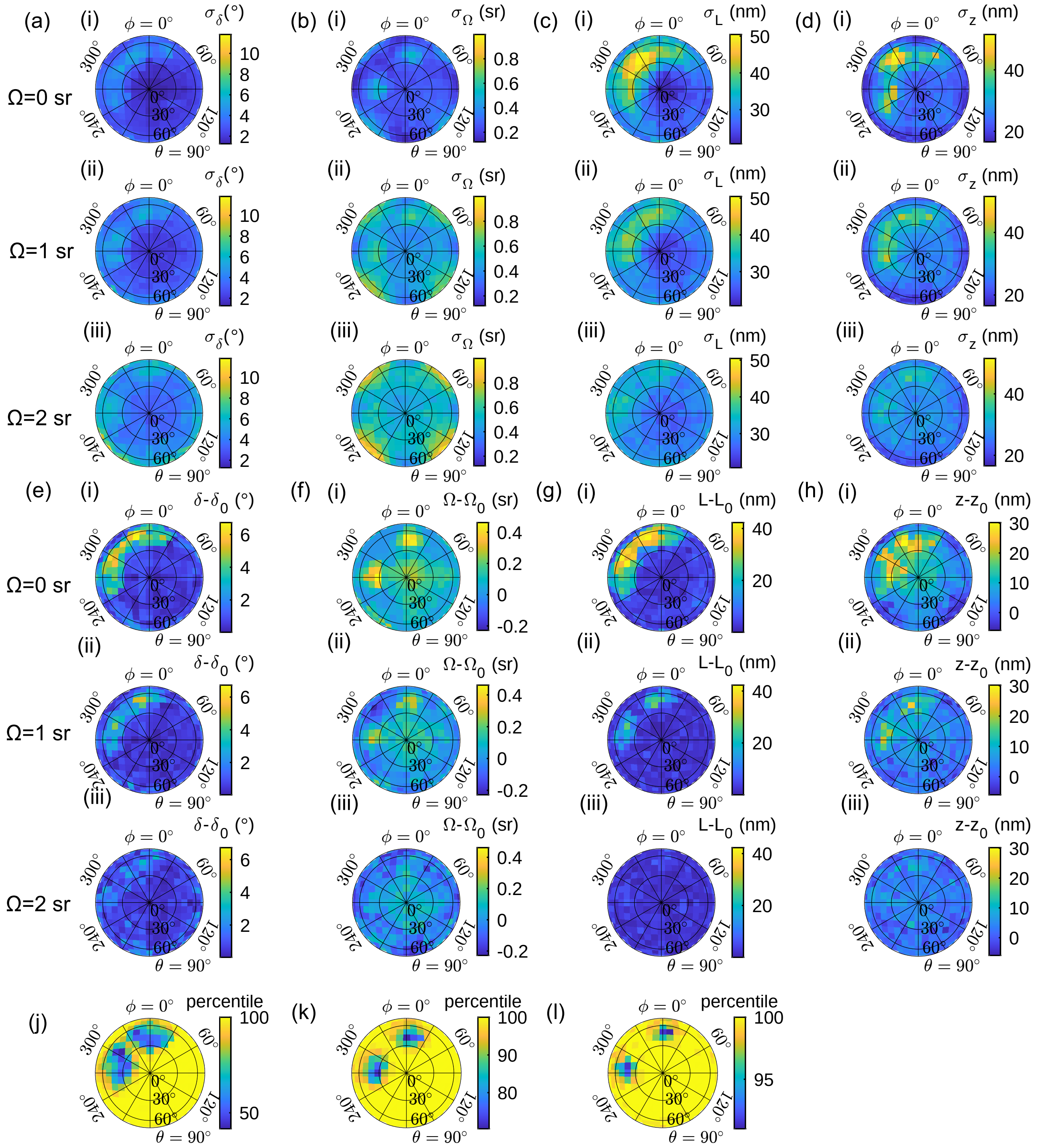

FIG. S12. Precision and accuracy of estimating 3D orientation and 3D position for emitters at $h = 700$ nm with a nominal focal plane placed at $z = -580$ nm after filtering biased estimates. Estimates >90 nm away from the ground truth 2D position $(x_0, y_0)$ are removed. (a-d) Precision of measuring 3D orientation (a) $\sigma_\delta$ (deg), (b) $\sigma_\Omega$ (sr), and 3D location (c) $\sigma_L$ (nm), (d) $\sigma_z$ (nm) at various orientations ($\Omega \in \{0, 1, 2\}$ sr, $\theta \in [0°, 90°]$, $\phi \in [0°, 360°)$) after filtering. (e-h) Accuracy of measuring 3D orientation (e) $\delta - \delta_0$ (deg), (f) $\Omega - \Omega_0$ (sr), and 3D location (g) $L - L_0$ (nm), (h) $z - z_0$ (nm). (j-l) Percentage of estimates passing the filter for (j) $\Omega = 0$ sr, (k) $\Omega = 1$ sr, and (l) $\Omega = 2$ sr. At each orientation, 200 independent images were generated for emitter within water located at $h = 700$ nm with 2500 signal photons and 3 background photons per pixel. The 3D orientation and 3D position are estimated as described in Section II.

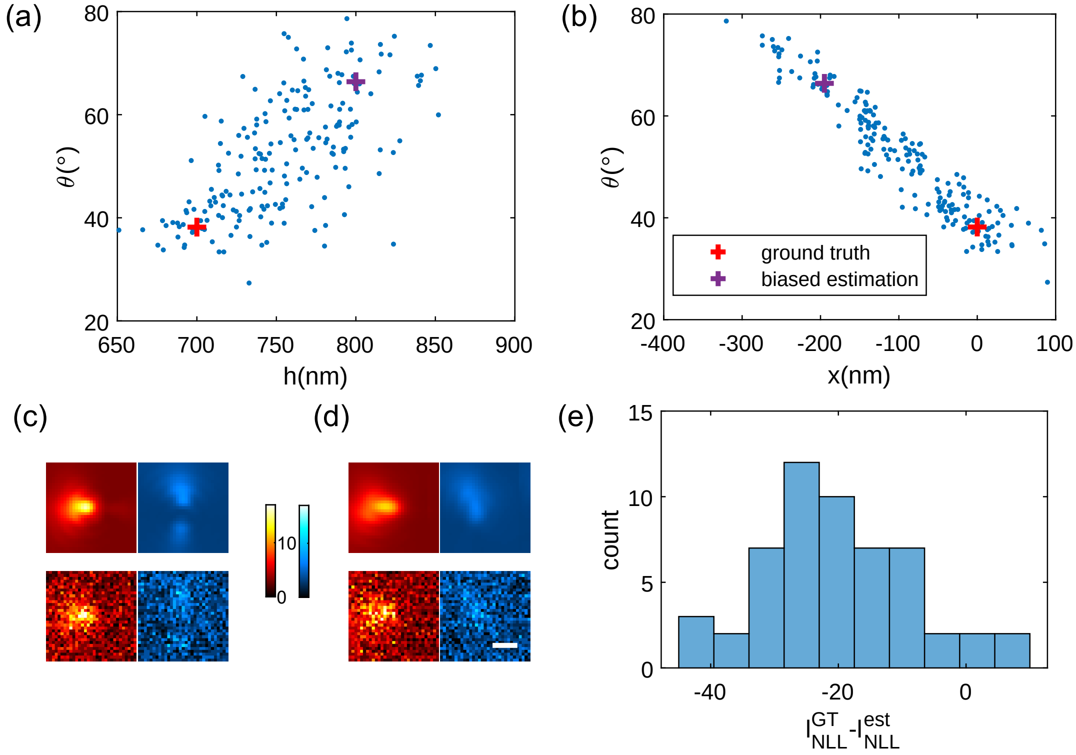

FIG. S13. Correlation between measurements of 3D orientation and 3D position for an emitter in water with a nominal focal plane placed at $z = -580$ nm. The emitter here is located at $(x_0, y_0, h_0) = (0, 0, 700)$ nm with 3D orientation $(\theta_0, \phi_0, \Omega_0) = (38°, 162°, 0)$. The algorithm in Section II is used to fit 200 independently generated noisy images. (a) Axial position estimation $h$ vs. polar angle estimation $\theta$. (b) Horizontal position estimation $x$ vs. polar angle estimation $\theta$. (c,d) (top) Noiseless PSF images and (bottom) images with Poisson shot noise for an emitter whose position and orientation match (c) the ground truth and (d) the biased estimate shown in (a,b). (Red) x-polarized, (blue) y-polarized. Colorbar: photons/pixel. Scalebar: 500 nm. (e) Comparing the value of the negative log-likelihood $l^{\mathrm{GT}}_{\mathrm{NLL}}$ of the ground truth (Eqn. S15) to the value of the negative log likelihood $l^{\mathrm{est}}_{\mathrm{NLL}}$ of the biased position and orientation (>90 nm away from the ground truth 2D position $(x_0, y_0)$). Negative values indicate that the ground truth position and orientation are more consistent with the noisy image than the biased estimate, i.e., $l^{\mathrm{GT}}_{\mathrm{NLL}} < l^{\mathrm{est}}_{\mathrm{NLL}}$, but our estimation algorithm (Sec. II) converged to the biased value instead.

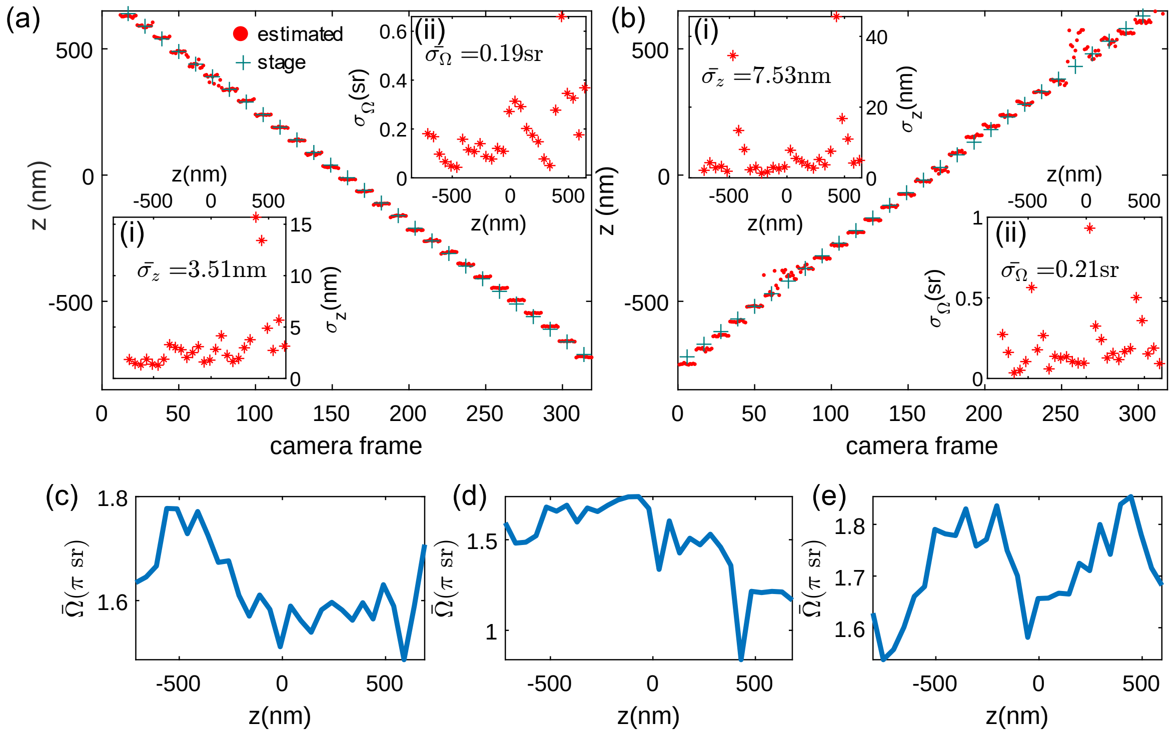

FIG. S14. Position and emission anisotropy measurements of two fluorescent beads. (a) Trajectory of defocus estimates $z$ for a bead scanned axially from $z = -790$ nm to $z = 610$ nm with a step size of 50 nm (11 camera frames per step). (b) Same as (a), but for another bead scanned in the opposite direction. Red dot: estimated axial distance $z$ between the bead and focal plane in each frame; green cross: expected stage position. Inset (i): Experimental axial precision $\sigma_z$ at each scanning plane (mean precision $\overline{\sigma_z}$ is (a) 3.51 nm and (b) 7.53 nm). Inset (ii): Experimental emission anisotropy precision $\sigma_\Omega$ at each scanning plane (average precision $\overline{\sigma_\Omega}$ is (a) 0.19 sr and (b) 0.21 sr). (c-e) Mean emission anisotropies of 3 fluorescent beads quantified as an effective wobble angle $\bar{\Omega}$ at each scanning plane: (c) bead in (a), (d) bead in (b), and (e) bead in Fig. 2(e).

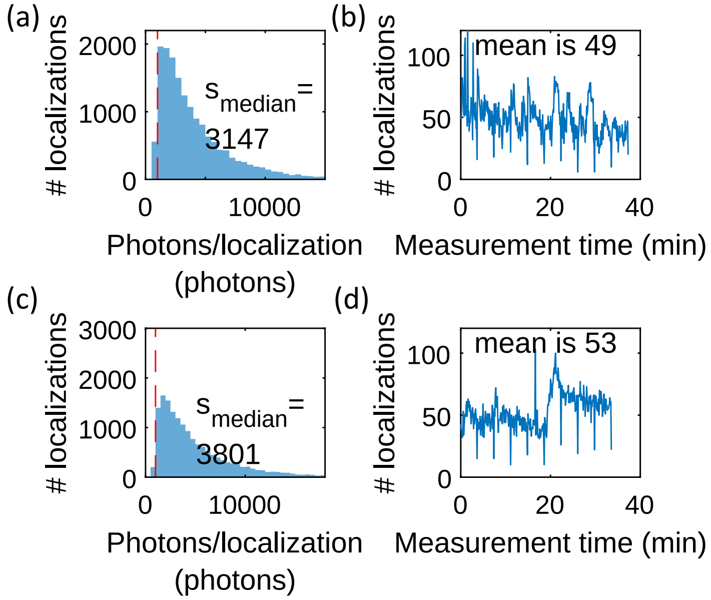

FIG. S15. Photophysics of Nile red (NR) blinking events. (a,c) Photons $s$ detected per localization of Nile red within spherical supported lipid bilayers (SLBs) consisting of (a) DPPC plus cholesterol and (c) DPPC only. The red dashed line represents $s = 1000$ photons. (b,d) Localization rate per 0.11 minutes for NR within (b) DPPC plus cholesterol and (d) DPPC-only SLBs.

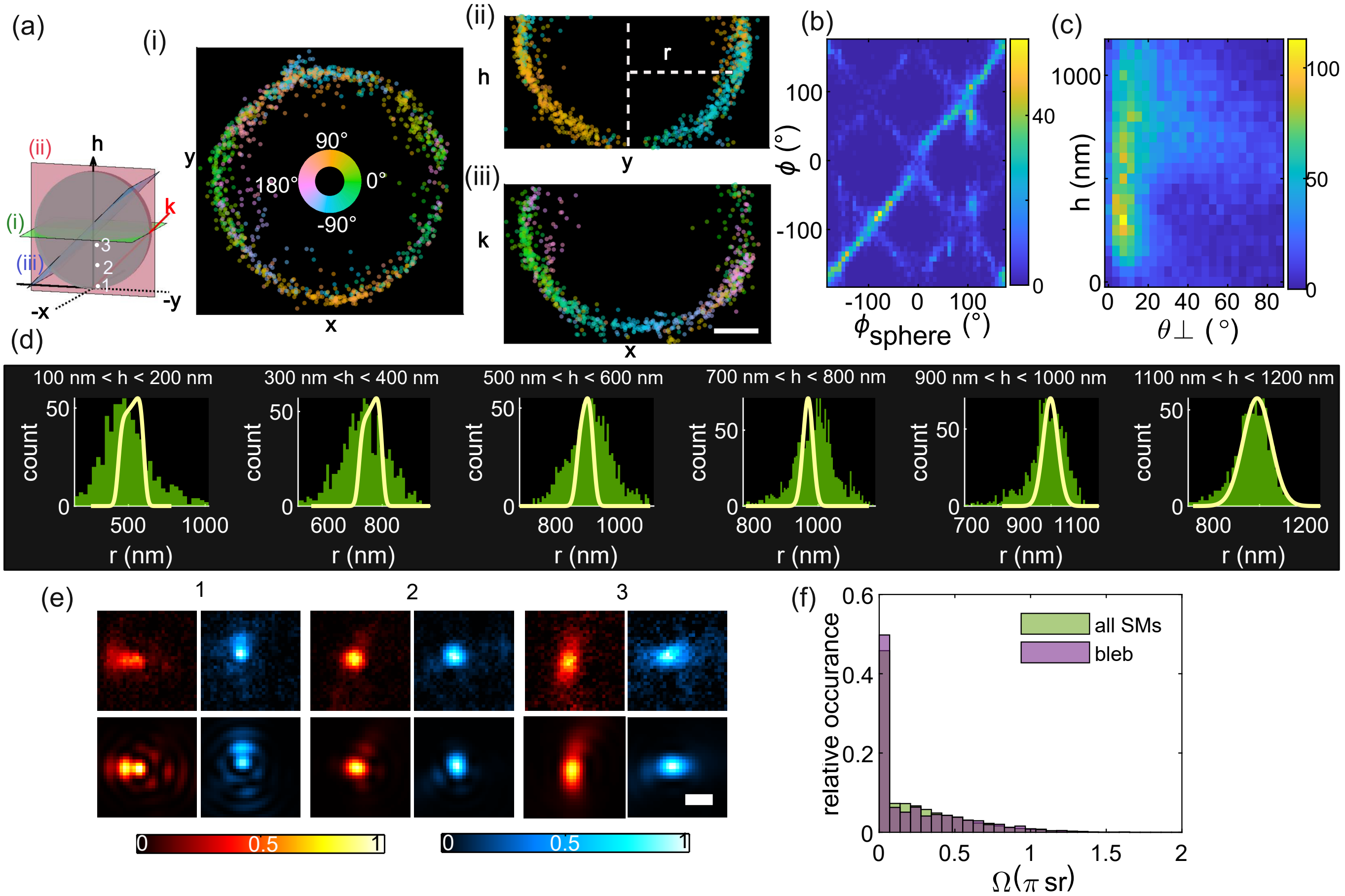

FIG. S16. SMOLM images of the 3D orientations and 3D locations of Nile red (NR) within a spherical supported lipid bilayer consisting of DPPC plus cholesterol. (a) Azimuthal angle $\phi$ of each NR molecule for the three cross-sections in inset: (i) x-y slice, (ii) y-h slice, and (iii) plane with y+h=1000 nm. Each slice has a thickness of 100 nm. Colorbar: $\phi$ (deg). Scalebar: 400 nm. (b) NR orientations $\phi$ relative to their positions on the sphere's surface $\phi_{\text{sphere}}$. (c) NR axial locations $h$ vs. orientations $\theta_{\perp}$. Colorbars: SM counts per bin. (d) NR lateral positions $r$ (see (a)(ii)) relative to the sphere's center within each $h$ slice compared to a model distribution (yellow lines, Eqn. S27), accounting for the curved spherical surface and pixOL's localization precision. (e, top row) Sum of all pixOL images of NR located within a $50 \times 50 \times 100$ nm$^3$ box at three locations in (a) inset: (1) $h = 150$ nm, (2) $h = 550$ nm, (3) $h = 950$ nm. (e, bottom row) Simulated pixOL images for emitters oriented perpendicular to the SLB and centered at the three dots locations in (a) inset with $\phi_{\text{sphere}} = 135°$. (Red) x-polarized, (blue) y-polarized. Colorbars: normalized intensity. Scalebar: 500 nm. (f) NR wobble $\Omega$ for (green) all localizations over the sphere and (purple) molecules within the membrane bleb in Fig. 3(e).

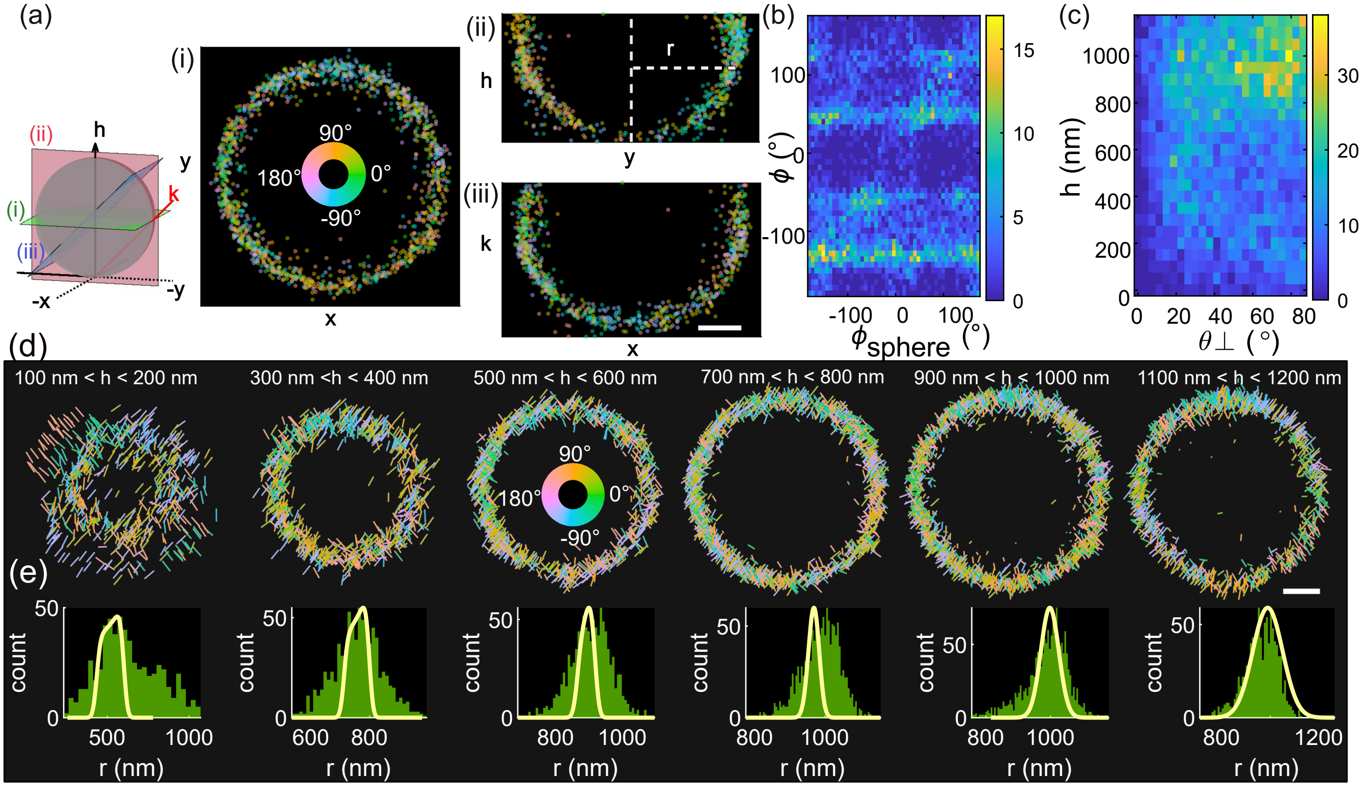

FIG. S17. SMOLM images of the 3D orientations and 3D locations of Nile red (NR) within a spherical supported lipid bilayer consisting of DPPC only. (a) Azimuthal angle $\phi$ of each NR molecule for three slices in inset: (i) x-y slice, (ii) y-h slice, and (iii) plane with y+h=1000 nm. Each slice has a thickness of 100 nm. Colorbar: $\phi$ (deg). (b) NR orientations $\phi$ relative to their positions on the sphere's surface $\phi_{\text{sphere}}$ for estimations with signal photons larger than 3000. (c) NR axial locations $h$ vs. orientations $\theta_\perp$ for estimations with signal photons larger than 3000. Colorbars: SM counts per bin. (d) x-y cross-sections of the bead depicting the 3D orientation ($\theta$,$\phi$) of each NR as a line segment. The length and direction of each line indicate the magnitude ($(\mu_x^2+\mu_y^2)^{1/2}$) and direction $\phi$ of azimuthal orientaion. Colors represent azimuthal orientation $\phi$. (e) NR lateral positions $r$ (see (a)(ii)) relative to the sphere's center within each $h$ slice compared to a model distribution (yellow lines, Eqn. S27), accounting for the curved spherical surface and pixOL's localization precision. Scalebars: 400 nm.

Movie S1 Demonstration of SM detection and position-orientation estimation using the pixOL microscope. (Top) Polarized images (red: x-polarized and blue: y-polarized) of Nile red are compared to (bottom) reconstructed images of the pixOL DSF using 3D orientations and 3D positions estimated by the algorithm described in Section II. Individual Nile red molecules are shown binding transiently to lipids (DPPC with cholesterol) within a spherical bilayer (Figs. 3 and S16). Colorbar: photons/pixel. Scale bar: 1 µm.

Movie S2 3D view of 6D SMOLM imaging of Nile red within spherical supported lipid bilayers consisting of (top) DPPC plus cholesterol (chol, Figs. 3 and S16) and (bottom) DPPC-only (Fig. S17). The position and orientation of each line depicts an emitter's 3D position and 3D orientation. Colors represent polar orientation $\theta$.

Movie S3 x-y cross-sections of 6D SMOLM imaging of Nile red within spherical supported lipid bilayers consisting of (top) DPPC plus cholesterol (chol, Figs. 3 and S16) and (bottom) DPPC only (Fig. S17) at various h-slices. The position and orientation of each line depicts an emitter's lateral position and 3D orientation. Colors represent azimuthal orientation $\phi$.
